# Regulation of hippocampal memory by mTORC1 in somatostatin interneuron

**DOI:** 10.1101/580670

**Authors:** J. Artinian, A. Jordan, A. Khlaifia, E. Honoré, A. La Fontaine, A-S. Racine, I. Laplante, J-C. Lacaille

## Abstract

Translational control of long-term synaptic plasticity via Mechanistic Target Of Rapamycin Complex 1 (mTORC1) is crucial for hippocampal learning and memory. The role of mTORC1 is well-characterized in excitatory principal cells but remains largely unaddressed in inhibitory interneurons. Here we used cell type-specific conditional knockout strategies to alter mTORC1 function selectively in somatostatin (SOM) inhibitory interneurons (SOM-INs). We found that up- and down-regulation of SOM-IN mTORC1 activity bi-directionally regulates contextual fear and spatial memory consolidation. Moreover, contextual fear learning induced a metabotropic glutamate receptor type 1 (mGluR1) mediated long-term potentiation (LTP) of excitatory input synapses onto hippocampal SOM-INs, that was dependent on mTORC1. Finally, the induction protocol for mTORC1-mediated late-LTP in SOM-INs regulated Schaffer collateral pathway LTP in pyramidal neurons. Thus, mTORC1 activity in somatostatin interneurons contributes to learning-induced persistent plasticity of their excitatory synaptic inputs and hippocampal memory consolidation, uncovering a role of mTORC1 in inhibitory circuits for memory.

## Introduction

Long-term synaptic plasticity is a prime candidate cellular substrate for learning and memory (*1–3*). In the hippocampus, long-term potentiation (LTP) at excitatory synapses of pyramidal cells is crucial for contextual as well as spatial learning and memory (*4*). In analogy to long-term memory consolidation, LTP in pyramidal cells displays an early phase, lasting minutes that is mediated by post-translational changes, and a late phase, lasting hours that requires transcription and translation (*3, 5*). Mechanistic Target of Rapamycin (mTOR) is a serine/threonine kinase that associates with two distinct protein complexes, mTOR Complex 1 and 2 (mTORC1 and 2), to control cell growth, proliferation and migration (*6*). mTORC1, of which Raptor is an essential component, plays a central role in cell growth by regulating protein synthesis and turnover, as well as lipid, nucleotide and glucose metabolism (*6*). In mature neurons, mTORC1 regulates protein synthesis during long-lasting synaptic plasticity and memory (*7, 8*). This role of mTORC1 in hippocampal memory was clearly established in principal cells (*7*).

In the hippocampus, excitatory neurons are regulated by highly heterogeneous inhibitory interneurons that display various morphological and physiological properties, as well as diverse patterns of connectivity and protein expression (*9*). Plasticity of hippocampal interneurons is also implicated in learning. Hippocampus-dependent learning is associated with structural plasticity of interneuron afferent connectivity (*10*) and modulation of their excitatory inputs (*11*). In addition, excitatory synapses onto hippocampal interneurons demonstrate multiple forms of LTP (*12, 13*). However, whether mTORC1 plays a role in these learning-related changes in inhibitory interneurons remains mostly unknown.

In the hippocampus, somatostatin (SOM) expressing interneurons (SOM-INs) are a subgroup of GABAergic interneurons that, in CA1, receive excitation from local principal pyramidal cells and provide feedback dendritic inhibition (*9*). They include the so-called Oriens-Lacunosum/Moleculare (O-LM) cells, bistratified cells and long-range projecting cells (*9*). SOM-INs regulate pyramidal cell synaptic integration (*14*), control their rate and burst firing (*15*), modulate their synaptic plasticity (*16*) and support contextual fear learning (*17*). Excitatory synapses onto SOM-INs express a form of LTP dependent on type 1a metabotropic glutamate receptors (mGluR1a) (*18, 19*). Additionally, a late form of mGluR1a-dependent LTP is present in O-LM interneurons in slice cultures (*8*) that lasts 24 hours and is dependent on mTORC1-mediated translation (*8, 20*). Thus SOM-INs represent an interesting subpopulation of interneurons to investigate mTORC1 function.

To determine if mTORC1 activity in SOM-IN plays a role in hippocampal memory, we used conditional knock-out mouse strategies to bi-directionally manipulate mTORC1 activity selectively in SOM-INs by targeting an essential component of mTORC1, the Regulatory-Associated Protein of mTOR (Raptor), or a repressor of mTORC1, the Tuberous Sclerosis Complex 1 (TSC1). We found that mTORC1 activity in somatostatin interneurons contributes to hippocampus-dependent memory formation and to learning-induced persistent plasticity at somatostatin interneuron input synapses, uncovering an important function of mTORC1 activity in inhibitory local circuits for memory consolidation.

## Results

We first established that mGluR1a- and mTORC1-dependent late-LTP (*8, 20*) occurs at excitatory synapses onto CA1 SOM-INs in mice expressing EYFP under the control of the SOM promoter (*Sst*^ires-Cre^;*Rosa26*^lsl-EYFP^). As previously reported for mature hippocampus (*19*), the vast majority of CA1 EYFP-expressing interneurons in slice cultures from *Sst*^ires-Cre^;^*Rosa26*lsl-EYFP^ mice were immuno-positive for SOM and located in stratum oriens (Fig. S1A). Whole-cell recordings from CA1 EYFP-expressing SOM-INs revealed that EPSCs were potentiated at 24h after repeated chemical mGluR1 stimulation (Fig. S1B). EPSC amplitude and potency were increased, and paired-pulse ratio was decreased, compared to sham treated slices (Fig. S1C, D and G). EPSC potentiation was prevented by the mGluR1a antagonist LY367385 (Fig. S1E and G) or the mTOR inhibitor PP242 (Fig S1F and G). Thus, persistent late-LTP occurs at synapses onto SOM-INs and is dependent on mGluR1a and mTORC1 signaling.

### Cell-specific Rptor deletion prevents mTORC1 signaling in SOM-INs

mTORC1 is a key regulator of translation in long-term synaptic plasticity (*7, 8*). We investigated the role of mTORC1 in SOM-INs in transgenic mice with a cell-specific homozygous knock-out of *Rptor* in SOM-INs (*Sst*^ires-Cre^;*Rosa26*^lsl-EYFP^;*Rptor*^fl/fl^ mice, termed Som-Raptor-KO) and wild-type control mice (*Sst*^ires-Cre^;*Rosa26*^lsl-EYFP^;*Rptor*^WT/WT^, termed Som-Raptor-WT). First we characterized the loss of mTORC1 function in SOM-INs in these mutant mice. The number of CA1 EYFP-expressing SOM-INs immuno-positive for Raptor was drastically reduced in Som-Raptor-KO mice compared to -WT, confirming a Raptor expression deficit in SOM-INs (WT: n=7, KO: n=8; Mann-Whitney test, *P*=0.0013; Fig. 1A). Next we confirmed the cell-type selectivity of *Rptor* deletion. Western blots assays of Raptor in hippocampal lysates (Fig. 1B) and cultured slices (Fig. 1C) showed no difference in total hippocampal Raptor and p-S6 expression in lysates (Raptor: WT: n=6, KO: n=6; *t*_10_=-0.8, *P*=0.44; p-S6S235/236: WT: n=3, KO: n=3; *t*_4_=-0.3, *P*=0.81) and cultured slices (Raptor: WT: n=2, KO: n=2; Mann-Whitney, *P*=1; p-S6S235/236: WT: n=3, KO: n=3; *t*_2_=-0.8, *P*=0.49) in Som-Raptor-KO and -WT mice, indicating that hippocampal principal neurons are unaffected by conditional deletion of *Rptor* in SOM-INs.

**Fig. 1:**
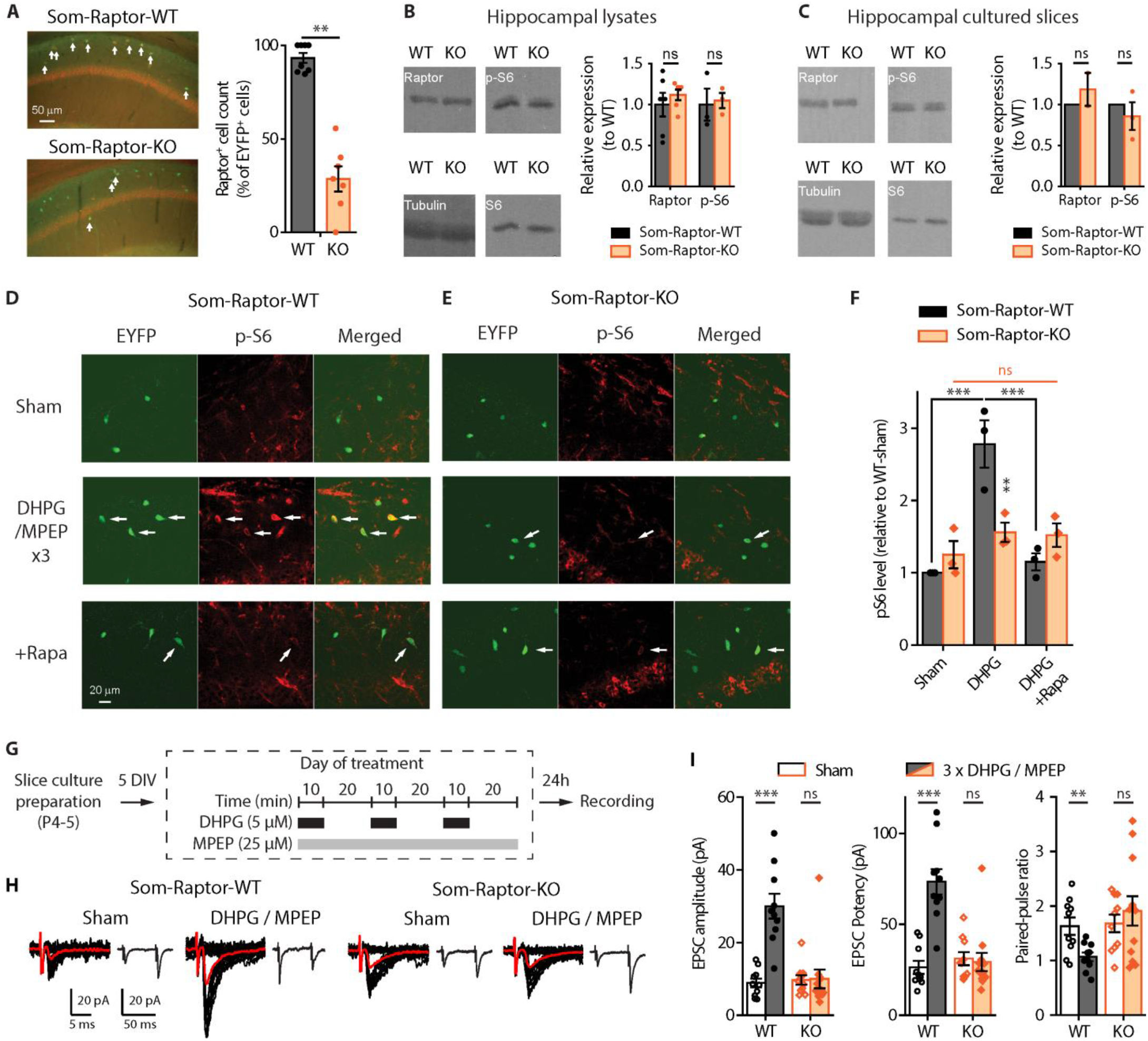
Cell-specific deficit in Raptor expression impairs mTORC1 signaling and late-LTP in SOM-INs of Som-Raptor-KO mice. (**A**) Left, Representative images of Raptor immunopositive (red) EYFP^+^ (green) CA1 SOM-INs (arrows, co-labeling) in Som-Raptor-KO relative to -WT mice. Right, Percentage of Raptor^+^ cells relative to EYFP^+^ cells. (B) Left, Representative Western blots of Raptor and phospho-S6^S235/236^ from hippocampal lysates. Right, Raptor (normalized to tubulin) and p-S6 (normalized to S6 and tubulin) levels (relative to Som-Raptor-WT mice). (**C**) Same as in (**B**) but in hippocampal cultured slices. (**D-E**) Representative confocal images illustrating EYFP^+^ CA1 SOM-INs (green), S6^S235/236^ phosphorylation (red) and co-labeling (merged) in Som-Raptor-WT (**D**) and -KO (**E**) mice, after sham, DHPG/MPEP and DHPG/MPEP in the presence of 1 μM rapamycin treatments. Arrows point to EYFP+ SOM-INs with p-S6 co-labeling. (**F**) Phosphorylated S6^S235/236^ level relative to sham treatment in Som-Raptor-WT mice. (**G**) Diagram of chemical late-LTP experimental protocol in cultured slices. (**H**) Representative EPSCs evoked by minimal stimulation in SOM-INs. Traces are superimposed 20 consecutive events (EPSCs + failures, black), average EPSC (including failures, red) and average of EPSC pairs (20 events) evoked by paired-pulse stimulation. (**I**) EPSC amplitude (including failures), EPSC potency (excluding failures) and paired-pulse ratio of SOM-INs 24h after DHPG/MPEP relative to sham treatment in Som-Raptor-WT and -KO mice.

We then verified impairment of mTORC1 signaling in SOM-INs using immunofluorescence (*8*). Repeated mGluR1 stimulation in hippocampal slices of Som-Raptor-WT mice increased phosphorylation of ribosomal S6 protein, a downstream effector of mTORC1, in SOM-INs, and this effect was prevented by the mTORC1 inhibitor rapamycin (Fig. 1D and F). Repeated mGluR1 stimulation failed to increase S6 phosphorylation in SOM-INs in Som-Raptor-KO mice, confirming the loss of mTORC1 signaling (Fig. 1E and F; two-way ANOVA, *F*_2,17(interaction)_=11.6, *P*=0.002; Bonferroni’s tests: WT-sham [n=3] vs. WT-DHPG [n=3], *P*=0.0002; WT-DHPG vs. WT-DHPG+rapamycin [n=3], *P*=0.0006; KO-sham [n=3] vs. KO-DHPG [n=3], *P*=1; KO-DHPG vs. KO-DHPG+rapamycin [n=3], *P*=1; WT-DHPG vs. KO-DHPG, *P*=0.008).

Next we assessed if mTORC1-mediated late-LTP was affected in SOM-INs. In slice cultures of Som-Raptor-WT mice, repeated mGluR1 stimulation (Fig. 1G), increased amplitude (sham: n=10, DHPG: n=10; t-test with Welch correction, *t*_11.2_=5.8, *P*=0.0001) and potency (Mann-Whitney, WT: *P*=0.0003), and reduced paired-pulse ratio (t-test with Welch correction, *t*_13.2_=3.2, *P*=0.007), of EPSCs in SOM-INs, relative to sham-treatment (Fig. 1H and I). Repeated mGluR1 stimulation failed to produce potentiation of EPSCs in Som-Raptor-KO mice (sham: n=12, DHPG: n=10; amplitude: Mann-Whitney tests, *P*=0.39; potency: *P*=0.34; paired-pulse ratio: t-test with Welch correction, *t*_20_=-0.7, *P*=0.48), indicating a block of persistent plasticity at excitatory synapses onto SOM-INs (Fig. 1H and I). SOM-INs basal excitatory synaptic transmission was intact as EPSCs were similar in Som-Raptor-KO and -WT mice in sham-treated conditions. Heterozygous Som-Raptor-KO mice failed to show a deficit in late-LTP (Fig. S1H). Together, these results show that conditional *Rptor* knockout in SOM-INs results in deficient mTORC1-mediated signaling and synaptic plasticity in SOM-INs.

As control, we verified that the numbers of CA1 SOM-INs were unaffected in Som-Raptor-KO relative to -WT mice (Fig. S2A and B), indicating that *Rptor* deletion driven by the SOM promoter did not alter SOM-INs proliferation and migration. In addition, because mTORC1 is a key regulator of protein synthesis implicated in cell growth (*6*), we used whole-cell current-clamp recordings and reconstruction of biocytin-filled SOM-INs in acute slices to show that general somatic and dendritic morphology, as well as basic membrane properties of CA1 SOM-INs were not deficient in Som-Raptor-KO mice (Fig. S2C to F and Fig. S3). These results indicate deficits in mTORC1 signaling and synaptic plasticity, with relatively intact morphological and membrane properties, in SOM-INs of conditional KO mice.

### SOM-IN specific Rptor deletion impairs hippocampus-dependent long-term memory

mTORC1 is a key regulator of translation in hippocampal long-term memory (*7*). Since hippocampal SOM-INs have been proposed to support contextual fear memories (*17, 21*), we examined the consequences of impairment of SOM-IN mTORC1 function in hippocampus-dependent memory tasks. We first verified in the open-field test that Som-Raptor-KO mice showed normal anxiety level and no impairment in locomotion, relative to -WT mice (Fig. S4). We next tested the mice in contextual fear learning and context discrimination (Fig. 2A). Som-Raptor-KO and -WT mice froze identically in response to footshocks, indicating normal anxiety and sensorimotor gating (WT: n= 55, KO: n=51; Mann-Whitney, *P*>0.05; Fig. 2B). In the short-term memory test (1h), Som-Raptor-KO and -WT mice showed similar freezing (WT: n= 11, KO: n=10; Mann-Whitney, *P*=0.70; Fig. 2C), indicating intact short-term contextual fear memory. However, Som-Raptor-KO mice displayed reduced freezing compared to -WT mice in the long-term memory test (24h, WT: n= 37, KO: n=34; *t*_69_=4.12, *P*=0.0001; Fig. 2C), revealing a long-term contextual fear memory deficit. Mice were also tested for context discrimination in one of two novel contexts varying in similarity with the training context (Fig. 2A). When exposed to a novel context, Som-Raptor-WT mice froze more in the similar (n=13) than in the distinct (n=13) context (Mann-Whitney, *P*=0.0021; Fig. 2D), manifesting context generalization and discrimination, respectively. Som-Raptor-KO mice showed reduced level of freezing in the similar novel context (n=14) compared to Som-Raptor-WT mice (Mann-Whitney, *P*=0.0009; Fig. 2D). Similar results were obtained using discrimination ratios to assess context discrimination normalized to the freezing level in the training context (two-way ANOVA, *F*_1,47(interaction)_=5.4, *P*=0.024; Tukey’s tests, WT-similar vs. WT-distinct: *P*=0.03; KO-similar vs. KO-distinct: *P*=0.96; WT-similar vs. KO-similar: *P*=0.003; WT-distinct vs. KO-distinct: *P*=0.50; Fig. 2D), indicating impairment in context generalization.

**Fig. 2:**
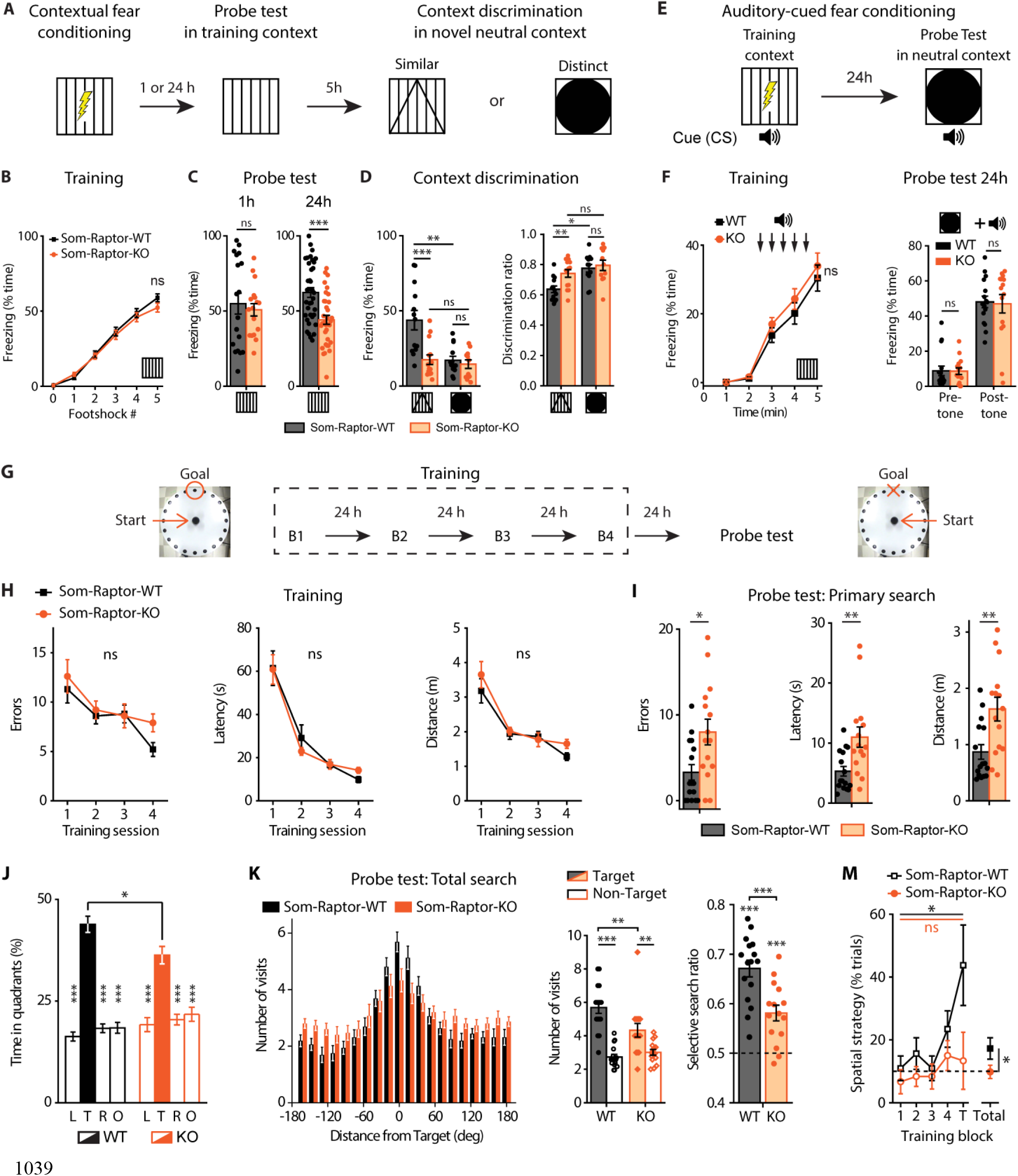
Som-Raptor-KO mice show impaired long-term contextual fear and spatial memory. (**A**) Diagram of the contextual fear conditioning protocol. (**B**) Percentage of time freezing after each footshock during the training session for Som-Raptor-WT and - KO mice (0: before the first footshock). (**C**) Percentage of time freezing during the probe tests at 1h (left) and at 24h (right). (**D**) Percentage of time freezing during the contextual discrimination test in a novel context (left) and discrimination ratio (right). (**E**) Diagram of the auditory-cued fear conditioning protocol. (**F**) Percentage of time freezing for Som-Raptor-WT and -KO mice during the training session (left) and during the probe test at 24h (right). (**G**) Diagram of the spatial learning protocol in the Barnes maze. (**H**) Performance in the Barnes maze during the training of Som-Raptor-WT and -KO mice. (**I**) Spatial memory performance during the primary search of the probe test. Errors, latency and distance before the first visit of the target. (**J**) Percentage of time spent in each quadrant of the maze during the total search period of the probe test. L: left, T: target, R: right, O: opposite quadrants. (**K**) Number of visits to all escape holes (left), number of visits expressed as target vs. average of all non-target holes (middle) and selective search ratios (right). (**M**) Percentage of trials using a spatial strategy over the training protocol. The dashed line represents chance (10%).

Fear conditioning strongly relies on amygdala function where SOM-INs play crucial roles (*22, 23*). We subjected mice to auditory-cued fear conditioning (Fig. 2E), a task that does not engage dorsal hippocampus, to assess if conditional *Rptor* deletion in SOM-INs affected amygdala function. Som-Raptor-KO and -WT mice showed similar levels of freezing in response to tone-shock presentations (WT: n= 16, KO: n=15; Mann-Whitney, *P*>0.05) and in the long-term memory probe test (Mann-Whitney, Pre-Tone and Post-Tone: *P*=0.57; Fig. 2F), indicating that mTORC1 function in SOM-INs is not required for dorsal hippocampus-independent cued fear memory.

To investigate further the behavioral role of mTORC1 in SOM-INs, we examined spatial reference memory in the Barnes maze, a hippocampus-dependent spatial learning and memory task (Fig. 2G). Som-Raptor-WT (n=16) and -KO (n=15) mice performed similarly during acquisition (errors: Friedman ANOVA, WT: *P*=0.0008; KO: *P*=0.036 and Mann-Whitney tests: S1: *P*=0.75, S2: *P*=0.87, S3: *P*=0.71, S4: *P*=0.08; latency: Friedman ANOVA, WT and KO: *P*<0.0001 and Mann-Whitney tests: S1: *P*=0.89, S2: *P*=0.78, S3: *P*=0.86, S4: *P*=0.013; distance: repeated measures ANOVA, *F*_3,29(session)_=16.2, *P*<0.0001; *F*_1,29(genotype)_=1.5, *P*=0.23; *F*_3,29(interaction)_=0.7, *P*=0.53), indicating intact spatial learning (Fig. 2H and fig. S5A). During the memory probe test, Som-Raptor-KO mice performed poorly with increased number of errors (Mann-Whitney, *P*=0.017), latency (*P*=0.007) and travelled distance (*P*=0.005) in the primary search for the target, indicating a deficit in long-term spatial memory at the single trial level, relative to -WT mice (Fig. 2I and fig. S5A). During the total search period, Som-Raptor-KO mice spent less time in the target quadrant than -WT mice (two-way ANOVA, *F*_3,116(interaction)_=5.4, *P*=0.002 and Tuckey’s tests, WT-target vs. KO-target: *P*=0.028; Fig. 2J and fig. S5A), indicating again an impairment in long-term spatial memory. In addition, Som-Raptor-KO mice visited less the target hole, relative to -WT mice, demonstrating a deficit in spatial memory precision (visits: Wilcoxon tests, WT: *P*=0.0005; KO: *P*=0.002 and Mann-Whitney test, WT-target vs. KO-target: *P*=0.007; selective search ratio: t-test against 0.5, WT: *t*_15_=10.2, *P*<0.0001; KO: *t*_14_=5.1, *P*=0.0002; WT vs. KO: *t*_29_=3.9, *P*=0.0006; Fig. 2K and fig. S5A). Mice solve the Barnes maze using thygmotactic/serial and/or hippocampus-dependent spatial strategies (*24, 25*). During the training protocol, Som-Raptor-WT mice rapidly manifested thygmotactic behaviors to solve the task, visiting holes in a serial manner but progressively adopted a spatial strategy, navigating directly to the target hole (Cochran test, *P*=0.015; Fig. 2M and fig. S5B). Som-Raptor-KO mice did not significantly use spatial strategy but solved the maze by relying mostly on thygmotactic or mixed strategies (Cochran test, *P*=0.67, WT: n=256 total trials, KO: n=240; z-score WT vs. KO: *P*=0.016; Fig. 2M and Fig. S5B), indicating an impairment in hippocampus-dependent spatial memory. Together, these results reveal that mTORC1 activity in SOM-INs is necessary for intact long-term spatial and contextual fear memory.

### Conditional Tsc1 knock-down augments mTORC1 signaling in SOM-INs

Tuberous Sclerosis Complex (TSC) is a key negative regulator of mTORC1 under basal conditions and upstream signaling pathways inhibit TSC to activate mTORC1 (*6*). We thus examined if *Tsc1* knock-down in SOM-INs would be sufficient to upregulate mTORC1 activity and promote hippocampal memory. Hence we used transgenic mice with a cell-specific heterozygous knock-out of *Tsc1* in SOM-INs (*Sst*^ires-Cre^;*Rosa26*^lsl-EYFP^;*Tsc1*^WT/fl^ mice, termed Som-TSC1-KO; and *Sst*^ires-Cre^;*Rosa26*^lsl-EYFP^;*Tsc1*^WT/WT^, termed Som-TSC1-WT). We first confirmed the cell-type selectivity of *Tsc1* knock-out. Western blots assays showed no difference in phosphorylated S6 protein in hippocampal lysates in Som-TSC1-KO and -WT mice (WT, n=12, KO: n=13; Mann-Whitney, *P*=0.25; Fig. 3B), indicating unaffected mTORC1 signaling in hippocampal principal neurons. Using immunofluorescence, basal level of phosphorylation of ribosomal S6 protein was increased in SOM-INs in slices of Som-TSC1-KO mice relative to -WT mice (WT, n=6, KO: n=6; *t*_5_=8.4, *P*=0.0004; Fig. 3C to E). Repeated mGluR1 (Fig. 3A) stimulation increased S6 phosphorylation in SOM-INs of Som-TSC1-WT relative to sham-treated (*t*_5_=3.2, *P*=0.024), but did not in -KO mice (t-test with Welch correction, *t*_6.1_=-0.8, *P*=0.46; Fig. 3C to E). Thus conditional Tsc1 knock-down increased basal mTORC1 activity and occluded mGluR1-induced mTORC1 activation in SOM-INs.

**Fig. 3:**
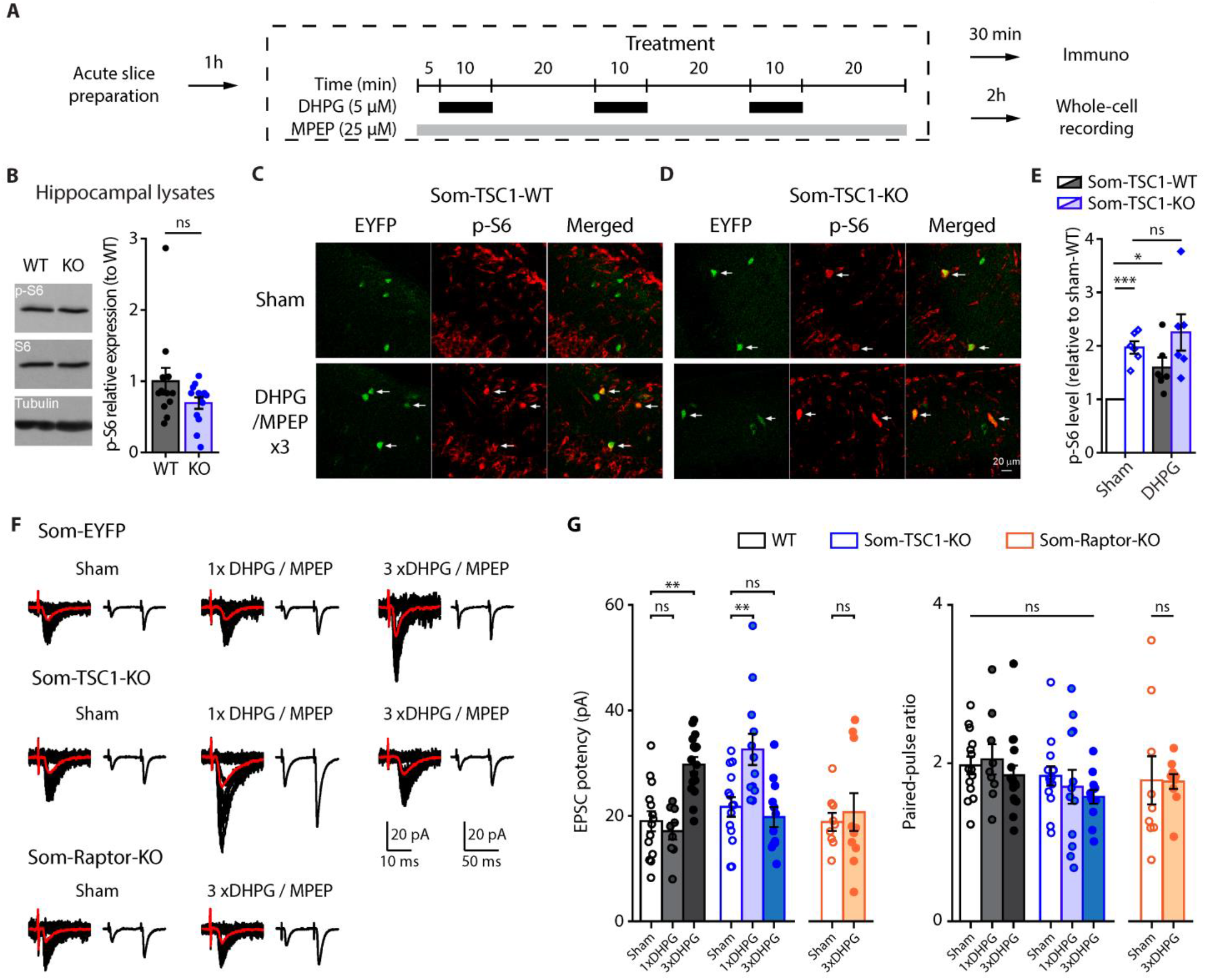
*Tsc1* knockdown increases mTORC1 signaling and facilitates mGluR1-mediated late-LTP in SOM-INs. (**A**) Diagram of chemical late-LTP experimental protocol in acute slices. (**B**) Left, Representative Western blots of phosphorylated S6^S235/236^ from hippocampal lysates. Right, p-S6^S235/236^ (normalized to S6 and tubulin) levels (relative to Som-TSC1-WT mice). (**C, D**) Representative confocal images illustrating EYFP+ CA1 SOM-INs (green), S6^S235/236^ phosphorylation (red) and co-labeling (merged) in Som-TSC1-WT (**C**) and -KO (**D**) mice, after sham and DHPG/MPEP treatments. Arrows point to EYFP+ SOM-INs with p-S6 co-labeling. (**E**) p-S6^S235/236^ level in the different groups relative to sham treatment in Som-TSC1-WT mice. (**F**) Representative EPSCs evoked by minimal stimulation in SOM-INs. Traces are superimposed 20 consecutive events (EPSCs + failures, black), average EPSC (including failures, orange) and average of EPSC pairs (20 events) evoked by paired-pulse stimulation. (**G**) EPSC potency (excluding failures) and paired-pulse ratio of SOM-INs 2h after sham, single and repeated DHPG/MPEP treatment in WT, Som-TSC1-KO and Som-Raptor-KO mice.

We then determined the effect of *Tsc1* knock-down on mTORC1-mediated synaptic plasticity in SOM-INs. First, current-clamp electrophysiological characterization of SOM-INs in slices of Som-TSC1-KO mice showed no major difference in basic membrane or firing properties, except a reduction in input resistance (Fig. S6). Next, we examined the effects of Tsc1 knock-down on chemically-induced late-LTP in SOM-INs (Fig. 3A). First we found that basal synaptic transmission was unaffected in the mutant mice. In sham treated slices, EPSC potency was unchanged in Som-TSC1-KO relative to -WT mice (Fig. 3F and G). In acute slices of WT mice, like in cultured slices, repeated mGluR1 stimulation was necessary to induce persistent potentiation of EPSC potency, relative to sham treatment (Fig. 3F and G). However in slices of Som-TSC1-KO mice, a single mGluR1 stimulation was sufficient, but repeated treatment failed, to induce persistent potentiation of EPSC potency (two-way ANOVA, *F*_2,71(interaction)_=19, *P*<0.0001, Bonferroni tests, WT-Sham [n=14] vs. WT-1xD [n=9]: *P*=0.99, WT-Sham vs. WT-3xD [n=16]: *P*=0.0014, TSC1KO-Sham [n=14] vs. TSC1KO-1xD [n=12]: *P*=0.0033, TSC1KO-Sham vs. TSC1KO-3xD [n=12]: *P*=0.98; Fig. 3F and G), indicating a facilitation of induction of late-LTP in SOM-INs with *Tsc1* knock-down. We verified that SOM-IN chemical late-LTP in acute slices was mTORC1-mediated and found that persistent potentiation of EPSC potency by repeated mGluR1 stimulation was blocked in SOM-INs of Som-Raptor-KO mice (WT: n=9, KO: n=10; t-test with Welch correction: *t*_12.7_=0.5, *P*=0.65; Fig. 3F and G). Chemical late-LTP in acute slices was associated with unchanged paired-pulse ratio (WT vs TSC1, two-way ANOVA, *F*_2,71(interaction)_=0.3, *P*=0.76, RaptorKO-Sham vs RaptorKO-3xD: t-test with Welch correction: *t*_9.5_=0.05, *P*=0.96; Fig. 3F and G), indicating some difference in mechanism with cultured slices. Together, these results suggest that conditional mTORC1 upregulation in SOM-INs results in a lower threshold for induction of persistent synaptic plasticity in SOM-INs, and in an impairment of plasticity elicited by a normal induction paradigm.

### Conditional Tsc1 knock-down in SOM-INs augments hippocampus-dependent longterm memory

Next we determined at the behavioral level the effects of *Tsc1* knock-down in SOM-INs. Som-TSC1-KO mice showed normal anxiety and slightly reduced locomotion in the open-field test (Fig. S7). Then we examined contextual fear memory and context discrimination (Fig. 4A). Som-TSC1-KO and -WT mice responded similarly to footshocks (WT: n=28, KO: n=31; Mann-Whitney tests, *P*>0.05; Fig. 4B), indicating intact sensorimotor gating. One hour after conditioning, Som-TSC1-KO and -WT mice showed similar freezing (WT: n=12, KO: n=13; *t*_23_=-0.76, *P*=0.46; Fig. 4C), indicating intact short-term contextual fear memory. However, in the 24h memory probe test, Som-TSC1-KO mice displayed increased freezing compared to -WT mice (WT: n=16, KO: n=18; t-test with Welch correction, *t*_22.2_=-3.08, *P*=0.005; Fig. 4C), revealing an augmentation of long-term contextual fear memory. When exposed to a novel distinct context, Som-TSC1-KO mice showed higher level of freezing (WT: n=8, KO: n=8; *t*_16_=3.57, *P*=0.0027) and lower discrimination ratio (*t*_16_=3.55, *P*=0.003) relative to -WT mice, indicating impairment in context discrimination (Fig. 4D). These results suggest that increased mTORC1 activity in SOM-INs is sufficient to augment long-term contextual fear memory and generalization. Next, we examined whether amygdala function was affected. Som-TSC1-KO mice were subjected to auditory-cued fear conditioning (Fig. 4E) and showed no difference in freezing during the training (WT: n=11, KO: n=13; Mann-Whitney tests, *P*>0.05; Fig. 4F) or 24h memory probe test (Mann-Whitney, Pre-Tone: *P*=0.16; Post-Tone: *P*=0.56; Fig. 4G), indicating that upregulated mTORC1 activity in SOM-INs does not alter this dorsal hippocampus-independent memory task.

**Fig. 4:**
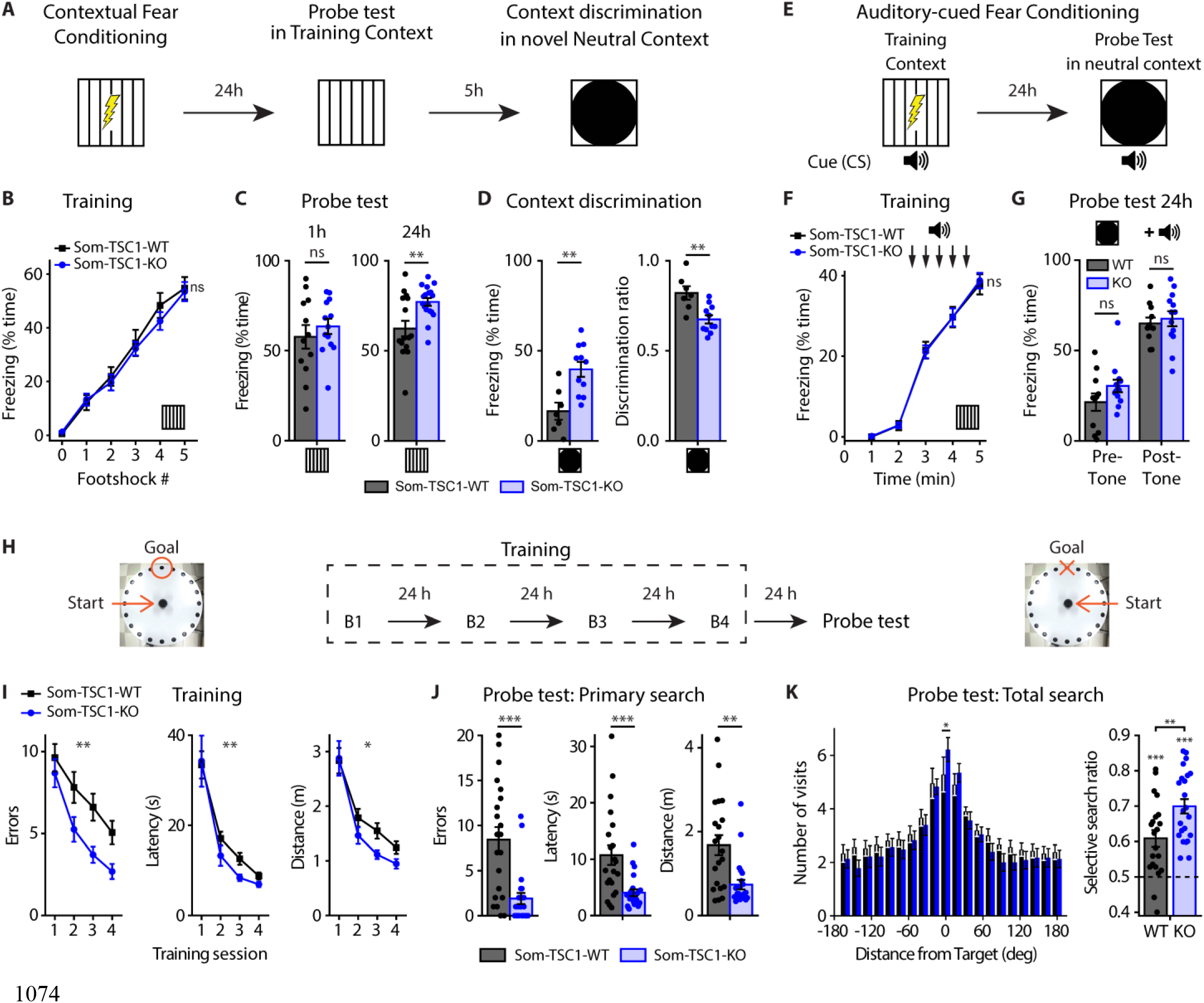
*Tsc1* deletion in SOM-INs increased long-term contextual fear and spatial memories but impaired context discrimination. (**A**) Diagram of the contextual fear conditioning protocol. (**B**) Percentage of time freezing after each footshock during the training session for Som-TSC1-WT and -KO mice (0: before the first footshock). (**C**) Percentage of time freezing during the probe tests at 1h and at 24h after conditioning. (**D**) Percentage of time freezing (left) and discrimination ratio (right) during the contextual discrimination test. (**E**) Diagram of the auditory-cued fear conditioning protocol. (**F, G**) Percentage of time freezing during the training session (**F**) and during the probe test 24h after (**G**) for Som-TSC1-WT and -KO mice. (**H**) Diagram of the spatial learning protocol in the Barnes maze. (**I**) Learning curves for Som-TSC1-WT and -KO mice during the training session. (**J**) Memory performance during the primary search of the probe test. (**K**) Number of visits to all escape holes (left) and selective search ratio (right) during the total search of the probe test.

Finally, we investigated spatial memory performance of Som-TSC1-KO mice in the Barnes maze (Fig. 4H). Som-TSC1-KO mice showed fewer errors, shorter latency and travelled distance to target compared to WT during the acquisition (WT: n=21, KO: n=23; errors: repeated two-way ANOVA, *F*_1,42(genotype)_=9.47, *P*=0.004; latency: Friedman ANOVA, WT and KO: *P*<0.0001 and Mann-Whitney tests, S1: *P*=0.26, S2: *P*=0.012, S3: *P*=0.0098, S4: *P*=0.048; distance: repeated two-way ANOVA, *F*_1,42(genotype)_=6.09, *P*=0.018; Fig. 4I and Fig. S8) and the long-term memory test (Mann-Whitney tests, errors: *P*=0.0003, latency: *P*=0.0009; distance: *P*=0.002; Fig. 4J and Fig. S8), demonstrating better spatial learning and memory performance. During the total search period, Som-TSC1-KO mice visited more the target hole (Mann-Whitney, *P*=0.047) and showed a higher selective search ratio than the -WT mice (t-test against 0.5, WT: *t*_20_=4.61, *P*=0.0002; KO: *t*_22_=9.76, *P*<0.0001; WT vs. KO: *t*_42_=-2.91, *P*=0.006; Fig. 4K and Fig. S8), indicating a better selective search of the target in Som-TSC1-KO mice.

These changes observed after *Tsc1* knock-down are the converse of changes found after *Rptor* deletion. Knock-down of *Tsc1* in SOM-INs upregulated mTORC1 activity and promoted hippocampal learning and memory. Thus mTORC1 activity level in SOM-INs appears to determine the level and contextual precision of hippocampal memories.

### Contextual fear learning induces mTORC1-mediated plasticity at SOM-IN excitatory synapses

Next we examined whether the effects of conditional genetic manipulations of mTORC1 on hippocampus-dependent memory and SOM-IN synaptic plasticity could be related. We tested if contextual fear learning produces potentiation at SOM-INs excitatory synapses via mTORC1 using ex vivo whole-cell recordings in acute slices 24h after training (Fig. 5A). SOM-INs from conditioned Som-Raptor-WT mice showed increased spontaneous EPSC frequency (naive: n=15; CFC: n=13; Mann-Whitney, *P*=0.015) and amplitude (Fig. 5B and C), increased minimally-evoked EPSC potency (naive: n=11; CFC: n=14; Mann-Whitney, *P*=0.0004) associated with decreased minimal stimulation intensity and unchanged paired-pulse ratio (Fig. 5D and E), as well as potentiation of evoked EPSC input-output function (Fig. 5F and G), compared to naive mice. Thus, conditioning induces potentiation of excitatory synapses onto SOM-INs lasting 24h. The learning-induced potentiation was mTORC1-dependent. At 24h post-training, SOM-INs of Som-Raptor-KO mice failed to show any potentiation of EPSCs relative to naive mice (spontaneous EPSC amplitude: naive: n=11; CFC: n=14; two-way ANOVA, *F*_1,49(interaction)_=5.1, *P*=0.004; Tukey’s tests, WT-naive vs. WT-CFC, *P*=0.007; KO-naive vs. KO-CFC, *P*=0.69; WT-naive vs. KO-naive: *P*=1; minimally-evoked EPSC potency: KO-naive [n=10] vs. KO-CFC [n=12]: *P*=0.87; paired-pulse ratio: two-way ANOVA, *F*_1,43(interaction)_=1.6, *P*=0.22; minimal stimulation intensity: two-way ANOVA, *F*_1,42(interaction)_=5.3, *P*=0.026; Tukey’s tests, WT-naive vs. WT-CFC: *P*=0.005; KO-naive vs. KO-CFC, *P*=0.99; WT-naive vs. KO-naive: *P*=0.31; input-output gain: two-way ANOVA, *F*_1,34(interaction)_=8.8, *P*=0.006; Tukey’s tests, WT-naive [n=9] vs. WT-CFC [n=10]: *P*=0.0001, KO-naive [n=9] vs. KO-CFC [n=10], *P*=0.89; WT-naive vs. KO-naive: *P*=0.99; Fig. 5B to G). SOM-INs from naive Som-Raptor-WT and -KO mice showed similar spontaneous and evoked EPSCs (Mann-Whitney, spontaneous EPSC frequency: *P*=0.32; minimally-evoked EPSC potency: *P*=0.25; Fig. 5B to G), indicating unchanged basal synaptic transmission. Furthermore, administration of the mGluR1 antagonist JNJ16259685 to Som-Raptor-WT mice 30 min before conditioning prevented the learning-induced potentiation (spontaneous EPSC frequency: vehicle [VEH]: n=15, JNJ: n=14; Mann-Whitney, *P*=0.014; amplitude: *t*_27_=2.13, *P*=0.0424; minimally-evoked EPSC potency: VEH: n=14, JNJ: n=12; *t*_24_=5.31, *P*<0.0001; minimal stimulation intensity: *t*_24_=-4.32, *P*=0.0002; paired-pulse ratio: *t*_24_=-0.61, *P*=0.55; input-output gain: VEH: n=12, JNJ: n=11; Mann-Whitney, *P*=0.0092; Fig. 5H). These results indicate that contextual fear learning induces mGluR1- and mTORC1-mediated persistent potentiation at SOM-INs excitatory synapses, suggesting that the effects of conditional genetic manipulations of mTORC1 on hippocampus-dependent memory and SOM-IN synaptic plasticity may be linked.

**Fig. 5:**
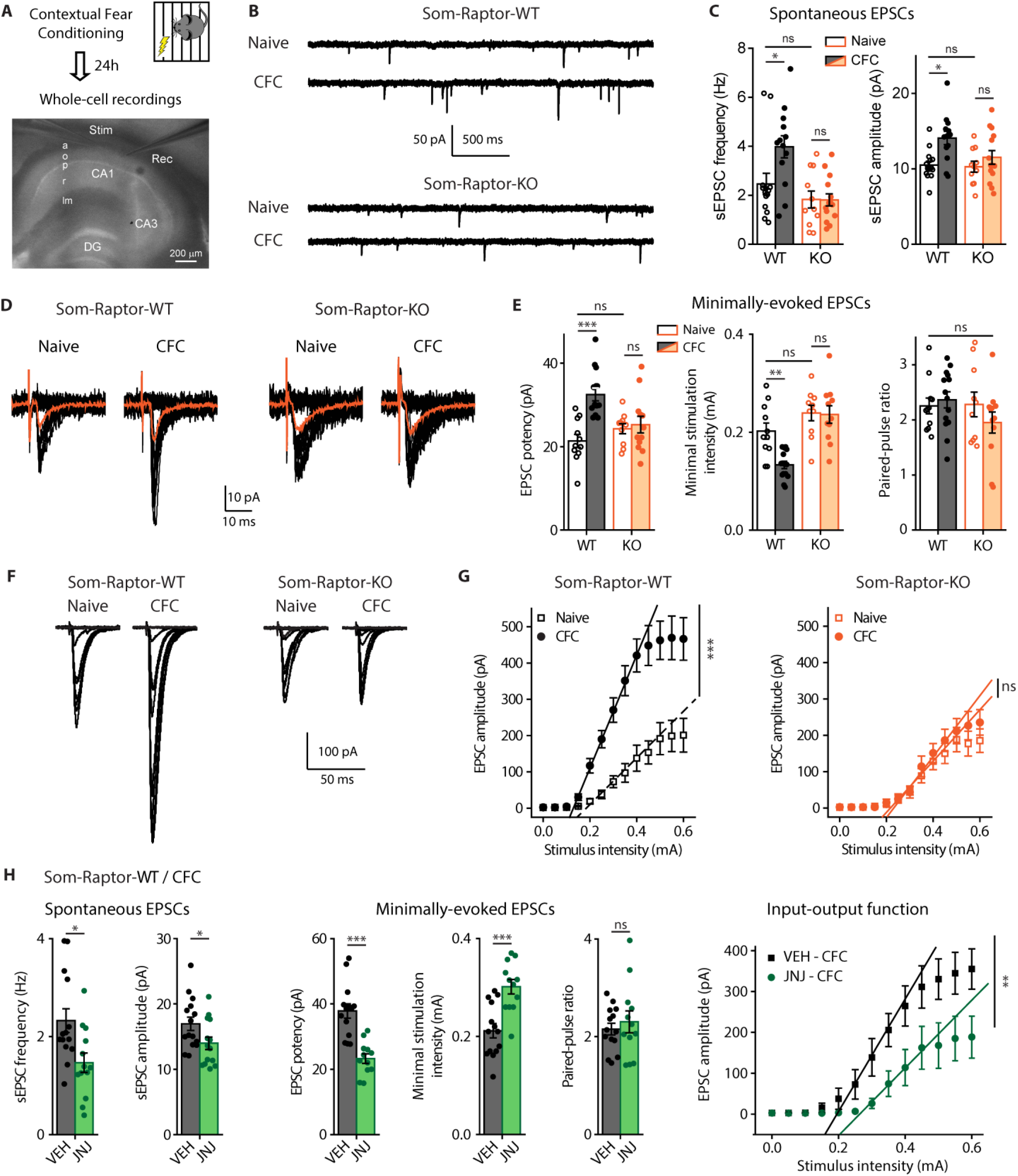
Contextual fear learning induces mGluR1- and mTORC1-mediated persistent LTP at excitatory synapses onto SOM-INs. (**A**) Diagram of the experimental protocol in acute slices. a: alveus, o: oriens, p: pyramidale, r: radiatum, lm: lacunosum-moleculare, stim: stimulation pipette, rec: recording pipette. (**B**) Representative traces of spontaneous synaptic activity in SOM-INs from naive and trained Som-Raptor-WT and -KO mice. (**C**) Spontaneous EPSC frequency and amplitude. (**D**) Representative traces of EPSCs evoked by minimal stimulation in the different conditions. (**E**) Minimally-evoked EPSC potency, paired-pulse ratio and minimal stimulation intensity. (**F**) Representative traces of evoked input-output function in SOM-INs in the different conditions. (**G**) Input-output gain of SOM-INs as the slope of individual linear regressions. (**H**) Synaptic properties of SOM-INs from CFC-trained Som-Raptor-WT mice treated with mGluR1 antagonist JNJ16259685 relative to vehicle (VEH). Spontaneous EPSC frequency and amplitude, minimally-evoked EPSC potency, minimal stimulation intensity and paired-pulse ratio, and input-output gain.

### mTORC1-mediated late-LTP induction protocol in SOM-INs regulates SC-LTP in pyramidal cells

CA1 area is the output of the hippocampus and LTP at CA3-CA1 pyramidal cell synapses is considered a crucial phenomenon induced by learning and supporting memory (*1, 2*). O-LM cells are a major subtype of dendrite-projecting SOM-INs (*9*) that provide differential regulation of SC and temporo-ammonic pathways onto CA1 pyramidal cells (*16*): they inhibit distal dendrites and downregulate LTP in the temporo-ammonic pathway; whereas they contact inhibitory interneurons in stratum radiatum and upregulate LTP at SC inputs (SC-LTP) by disinhibition (*16, 19, 26, 27*). Consequently, induction of mGluR1a-mediated early-LTP at excitatory synapses onto SOM-INs upregulates SC-LTP of pyramidal cells, indicating that plasticity at SOM-INs input synapses regulates metaplasticity of the CA1 network (*19*). Therefore, we next examined if mTORC1-mediated plasticity at SOM-IN input synapses controlled in a longer-lasting manner their output and regulated metaplasticity of the SC CA1 network. First, we established that late-LTP was elicited at SOM-INs synapses using electrical stimulation in acute slices (Fig. 6A). Repeated TBS stimulation at the oriens/alveus border in slices of Som-Raptor-WT mice induced, at 2h post-induction, increases in EPSC amplitude (control: n=12; TBS: n=12; Mann-Whitney tests, *P*=0.004) and potency (*P*=0.023), and decreases in paired-pulse ratio (*P*=0.026), relative to unstimulated slices (Fig. 6B and C). In contrast, SOM-INs from Som-Raptor-KO mice failed to show potentiation of EPSCs (control: n=11; TBS: n=11; Mann-Whitney, amplitude: *P*=0.74; potency: *P*=0.65; paired-pulse ratio: *P*=0.6; Fig. 6B and C), indicating that late-LTP elicited by repeated TBS was mediated by mTORC1.

**Fig. 6:**
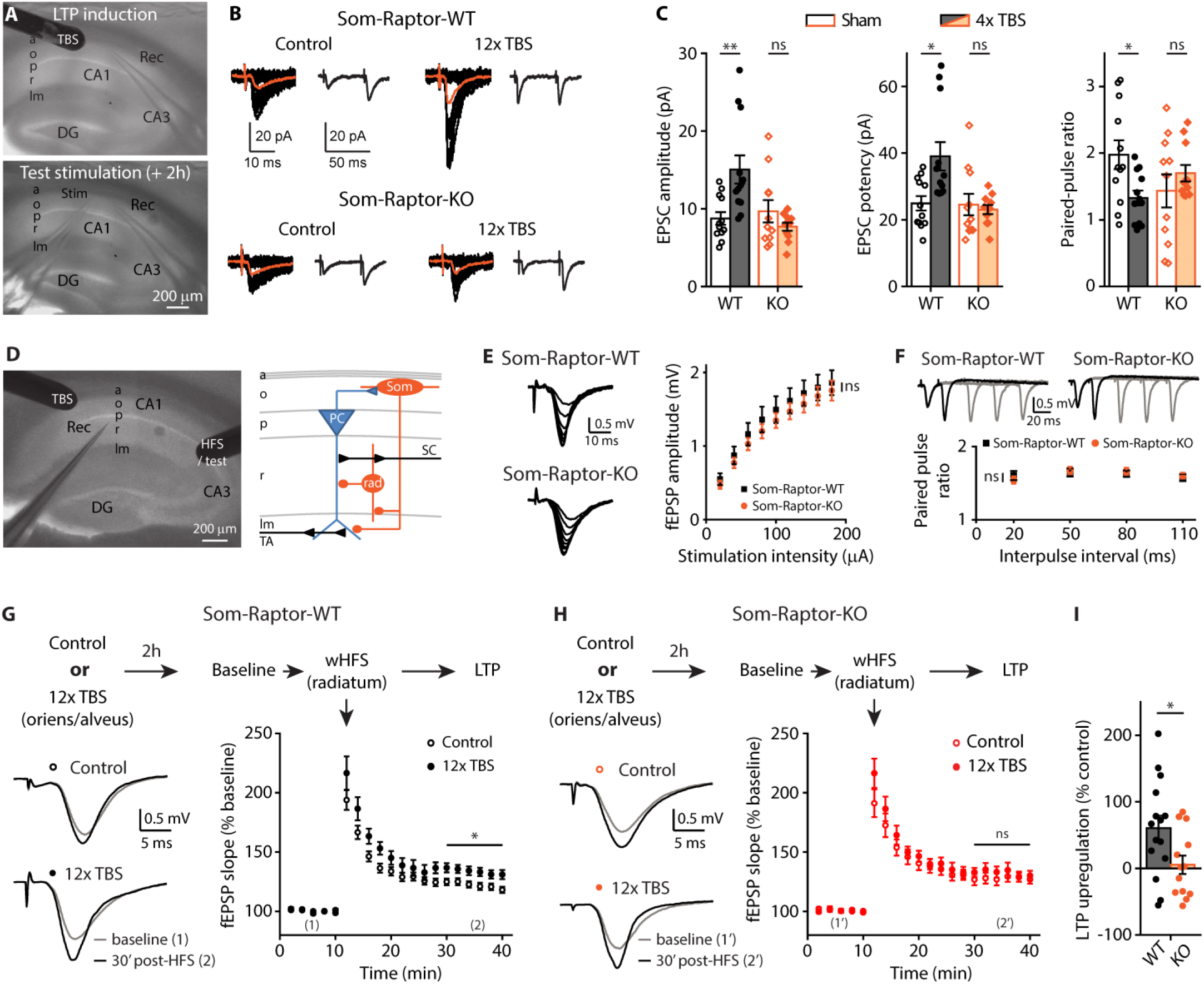
mTORC1-mediated late-LTP induction protocol in SOM-INs upregulates SC-LTP in principal cells. (**A**) Recording configuration for late-LTP induction and whole-cell recording 2h later. (**B**) Representative traces of EPSCs evoked by minimal stimulation in control condition and after repeated TBS stimulation, in Som-Raptor-WT and -KO mice. Traces are superimposed 20 consecutive events (EPSCs + failures, black), average EPSC (including failures, orange) and average of EPSC pairs (20 events) evoked by paired-pulse stimulation. (**C**) EPSC amplitude, potency (excluding failures) and paired-pulse ratio. (**D**) Recording configuration and simplified diagram of underlying CA1 network. SC: Schaffer collaterals, TA: temporo-ammonic pathway, PC: pyramidal cell, Som: SOM-INs, rad: radiatum interneuron. (**E**) Representative traces and amplitude of field EPSPs in response to incremental SC stimulation in Som-Raptor-WT and -KO mice. (**F**) Representative traces and ratio of field EPSPs amplitude in response to SC paired-pulse stimulations. (**G**) Top: Description of the stimulation and induction protocol. Left: representative traces of field EPSPs in response to SC stimulation 10 min before (1) and 30 min after weak HFS (wHFS, 100 Hz, 750 ms) stimulation (2) to induce SC-LTP, in control condition and 2h after the SOM-INs late-LTP induction protocol (repeated TBS stimulation) in Som-Raptor-WT mice. Right: Graph of LTP of field EPSP slope induced by wHFS. (**H**) Same as (**G**) in Som-Raptor-KO mice. (**I**) SC-LTP upregulation 2h after SOM-INs late-LTP induction protocol, normalized to the averaged control SC-LTP.

Having established a TBS protocol inducing mTORC1-mediated late-LTP in SOM-INs, we investigated its ability to control their output and upregulate SC-LTP of pyramidal cells in a persistent manner. We applied the repeated TBS induction protocol at the oriens/alveus border and two hours later applied weak high-frequency stimulation (wHFS) in the stratum radiatum to induce SC-LTP in pyramidal cells (Fig. 6D). SC-LTP magnitude was increased in slices of Som-Raptor-WT that received 2h previously the SOM-INs late-LTP induction protocol, relative to previously unstimulated slices (control: n=13; TBS: n=15; Mann-Whitney, *P*=0.03; Fig. 6G and I). The upregulation of SC-LTP by repeated TBS was prevented in slices from Som-Raptor-KO mice (control: n=9; TBS: n=13; KO: Mann-Whitney, *P*=0.89; WT vs. KO: *t*_26_=2.3, *P*=0.029; Fig. 6H and I), indicating that the upregulation of SC pathway plasticity required mTORC1-signaling, and thus likely mTORC1-mediated late-LTP, in SOM-INs. The block of SC-LTP regulation was not due to alterations in SC basal synaptic properties since input-output function (WT: n=9; KO: n=10; repeated two-way ANOVA, *F*_8,135(interaction)_=0.2, *P*=0.7; Fig. 6E) and paired-pulse facilitation (two-way ANOVA, *F*_3,76_=0.39, *P*=0.76; Fig. 6F) were similar in Som-Raptor-KO and -WT mice. These results show that repeated episodes of TBS induces long-lasting mTORC1-mediated plasticity at SOM-INs synapses that may result in long-lasting regulation of their output and CA1 network metaplasticity.

Together, our findings suggest that learning induces SOM-IN mTORC1 activity, resulting in persistent LTP at SOM-IN excitatory input synapses, and that this SOM-IN plasticity may in turn regulate CA1 SC synaptic plasticity. These SOM-IN mTORC1 cellular and circuit mechanisms could contribute to mTORC1 inhibitory circuit changes in contextual and spatial memory.

## Discussion

We used cell-type specific transgenic mouse approaches to reveal several novel insights about mTORC1 activity in SOM-INs for hippocampal function. At the behavioral level, we uncovered that loss of mTORC1 function in SOM-INs impaired contextual fear and spatial long-term memories, but spared sensory-motor gating, hippocampus-dependent short-term contextual memory and hippocampus-independent long-term auditory-cued fear memory. In contrast, upregulation of mTORC1 activity in SOM-INs by cell-specific conditional knockdown of *Tsc1* augmented spatial and contextual fear memories, and impaired discrimination. At the cellular level, bi-directional regulation of mTORC1 activity in SOM-INs differentially regulated mGluR1-mediated late-LTP at SOM-IN excitatory inputs. Moreover, contextual fear learning persistently increased afferent excitatory synaptic strength in SOM-INs via mGluR1 and mTORC1. At the network level, the SOM-IN late-LTP induction protocol upregulated metaplasticity of the SC pathway in pyramidal cells, in mTORC1-dependent manner. Our findings uncover a role of SOM-IN mTORC1 in learning-induced plasticity of their synapses and in hippocampal long-term memory consolidation.

Raptor is an essential component of mTORC1 which plays a central role in cell growth (*6*) in addition to regulation of protein synthesis during synaptic plasticity in mature neurons (*7*). We verified in control experiments that conditional recombination of *Rptor* in cells expressing Cre recombinase under control of the somatostatin promoter, resulted in effective loss of mTORC1 function sufficiently late not to impair development of SOM-INs. Som-Raptor-KO mice showed normal density of EYFP expressing SOM-INs in CA1 hippocampus, intact soma and dendritic morphology of individually labelled SOM-INs, as well as intact dendritic spine size and density. SOM-INs showed no change in basic membrane properties, except for a moderate increase of firing rate during sustained depolarization, as well as intact basal transmission and short-term plasticity of excitatory synaptic inputs. Thus the late-occurring conditional deletion of *Rptor* in SOM-Raptor-KO mice allowed us to examine mTORC1 function in mature SOM-INs. Similar observations were made with *Tsc1* knockdown that did not affect SOM-IN basic membrane properties, except a modest reduction of input resistance, nor basal transmission and short-term plasticity of excitatory synaptic inputs. *Rptor* deletion and *Tsc1* knockdown were not restricted in the hippocampus but occurred in all SOM-expressing cells and could possibly affect hippocampus-independent functions at the behavioral level. However, the intact anxiety, the moderate change in locomotion and the absence of freezing difference between WT and KO mice during CFC and short-term contextual fear memory indicate that hippocampus-independent functions such as sensorimotor gating and attention were spared in both conditional KO mice lines. Fear learning-induced synaptic potentiation in SOM-INs in the amygdala gates auditory-cued fear expression (*22, 23*). In the present study, conditional deletion of *Rptor* and *Tsc1* knockdown in SOM-INs did not affect dorsal hippocampus-independent long-term auditory-cued fear memory, indicating that learning-induced LTP in amygdala SOM-INs crucial for cued fear memory may not require mTORC1 mechanisms. Thus, learning-related SOM-INs synaptic plasticity mechanisms are heterogeneous with mTORC1 mechanisms prominent in hippocampal SOM-INs synaptic plasticity and showing some brain region-specificity.

The control of protein synthesis by mTORC1 in synaptic plasticity and memory consolidation is well-characterized in CA1 pyramidal cells (*7*). Our findings suggest that some aspects of mTORC1 function in synaptic plasticity are cell type-specific. We show that activation of mGluR1 in SOM-INs results in mTORC1-mediated late-LTP of excitatory synapses (*8*). However, activation of group I mGluRs in pyramidal cells was shown to induce mTORC1-mediated long-term depression of excitatory synapses (*28*). More recently, mTORC2 but not mTORC1 was associated with group I mGluR-mediated long-term depression (LTD) of CA1 pyramidal cells and novel object recognition (*29*). Thus, mGluR-mediated mTORC1, and perhaps even mTORC2, translational control likely regulates different mRNAs to achieve depression versus potentiation of synapses in pyramidal cells and SOM-INs, respectively. In addition, in other hippocampal interneurons which provides perisomatic inhibition of pyramidal cells, endocannabinoid-dependent long-term depression of their output synapses onto pyramidal cells requires presynaptic mTORC1-mediated protein synthesis to downregulate GABA release (*30*). The identification of specific mRNAs controlled by mTORC1 in different neurons classes will be important to clarify mTORC1 function in synaptic plasticity.

Cell type-specificity of mTORC1 function is consistent with findings that mTORC1 regulation of translation via one of its major effector, the translational repressor eukaryotic Initiation Factor 4E Binding Protein (eIF4E-BP), controls excitatory synaptic transmission differently in pyramidal cells and interneurons. In pyramidal cells, knock-out of eIF4E-BP results in enhanced basal excitatory synaptic transmission (*31*), as well as in a lower threshold for late-LTP induction and in impairment of late-LTP elicited by normal induction paradigms (*32*). In contrast, in hippocampal O-LM inhibitory interneurons, eIF4E-BP knockout does not affect basal excitatory transmission, lowers the threshold for late-LTP induction and does not affect late-LTP elicited by normal induction paradigm (*8*). Our results in SOM-INs of Som-TSC1-KO mice are largely similar to those in eIF4E-BP knockout. Conditional knockdown of *Tsc1* in SOM-INs increased mTORC1 activity as indicated by increase S6 phosphorylation in basal conditions. Moreover, the increased basal activity occluded mGluR1-induced mTORC1 activity in SOM-INs. Consistent with increased mTORC1 activity, the threshold for mGluR1-dependent late-LTP induction was lowered in SOM-INs of Som-TSC1-KO mice, with no change in basal excitatory transmission. Thus, upregulated mTORC1 activity, in combination with ongoing synaptic activity, is not sufficient to increase synaptic strength and mGluR1 activation is also required. Interestingly, repeated mGluR1 activation failed to induce late-LTP in SOM-INs of Som-TSC1-KO mice, suggesting the presence of a mechanism of homeostatic control downstream of mTORC1, possibly via S6 kinase/4E-BP2/eIF4E pathways, to prevent excessive or unselective translation of mRNAs (*32*).

We demonstrated that Som-TSC1-KO mice manifested increased contextual fear and spatial memories as well as altered context discrimination, suggesting that upregulating mTORC1 activity in SOM-INs is sufficient to alter hippocampal memory consolidation. These effects of *Tsc1* knockdown are converse to those of *Rptor* deletion and suggest that the level of mTORC1 activity in SOM-INs determines the level of persistent synaptic plasticity in SOM-INs and memory consolidation. Tuberous Sclerosis is a genetic disease due to mutation in TSC genes resulting in excessive mTORC1 activity and severe neurologic and psychiatric manifestations (*33*). Our findings in Som-TSC1-KO mice raise the interesting possibility that dysregulated mTORC1 activity and synaptic plasticity in SOM-INs contribute to cognitive impairments in Tuberous Sclerosis.

O-LM interneurons represent a major group of SOM-INs in CA1 hippocampus. They inhibit the distal temporo-amonic inputs of pyramidal cells in stratum lacunosum-moleculare (*9, 17*), and send axonal projections in stratum radiatum to inhibit interneurons that, in turn, inhibit SC inputs of pyramidal cells (*16*). O-LM interneurons differentially control plasticity of the two inputs to pyramidal cells: they inhibit and restrict LTP of temporo-amonic synapses; and they disinhibit and promote LTP of SC synapses (*16, 19, 26, 27*). Hebbian mGluR1a-mediated LTP at excitatory input synapses onto O-LM interneurons is translated into increased synaptically-driven action potential firing (*34*), providing a synaptic basis for increased inhibition of their downstream targets. Furthermore, induction of Hebbian mGluR1a-mediated LTP in SOM-INs upregulates for a period of minutes LTP in the SC pathway of pyramidal cells (*19*). Here we found that mTORC1-mediated late-LTP in SOM-INs upregulated for a period of hours LTP in the SC pathway. Interneurons expressing SOM in the CA1 hippocampus include not only O-LM cells but also bistratified cells and projection cells (*9*). In the present study, 90% of the biocytin labeled cells in our recordings were O-LM cells, clearly indicating a role of this interneuron subtype in SOM-IN mTORC1 actions in CA1. The development of new biological markers selective of other SOM interneuron subtypes will be necessary to assess their specific functional contributions.

We identified different physiological conditions in which mTORC1 activity is necessary for persistent enhancement of SOM-INs excitatory synapses: activation of mGluR1 in cultured (*8, 20*) and acute slices, stimulation of afferent fibers in acute slices, and contextual fear learning. Our results suggest that the persistent potentiation of excitatory synapses onto SOM-INs elicited in these conditions share some common mechanisms. Contextual fear conditioning may engage activity of CA1 pyramidal cell excitatory synapses and activation of local SOM-INs to induce Hebbian mGluR1-mediated late-LTP at these synapses (*8, 20*). Hence, our results are consistent with a model (Fig. S9) that hippocampal learning i) induces persistent mGluR1 and mTORC1-mediated late-LTP at CA1 SOM-INs synapses due to coincident activation of presynaptic local pyramidal cells and postsynaptic SOM-INs, ii) resulting in a long-lasting disinhibition of SC inputs to pyramidal cells due to increased inhibition of intercalated inhibitory interneurons, effectively leading to an upregulation of CA1 network metaplasticity by enhancing LTP at pyramidal cell SC synapses, and iii) causing improved consolidation of accurate hippocampal memory. In association with long-term depression of inhibition from cholecystokinin (CCK)-positive interneurons in stratum radiatum (*35*), and with the action of long-range inhibitory projections from entorhinal cortex (*36*), mTORC1-mediated LTP in SOM-INs may be a complementary mechanism for disinhibition of SC inputs in pyramidal cells (*27*). Indeed during spatial navigation, it was recently reported that, as the animal crosses a place field, CA1 place cells decrease their synaptic coupling with parvalbumin-expressing interneurons and increase it with SOM-INs, causing a switch of pyramidal cell inhibition from perisomatic/proximal dendritic to distal dendritic compartments, and allowing CA3 excitatory inputs to gain control over entorhinal excitatory inputs in driving pyramidal cell firing in a short-term fashion (*37*). Thus a crucial feature of mTORC1-mediated SOM-INs plasticity in spatial/contextual information encoding by CA1 pyramidal cells may be to promote internal representations by the hippocampal CA3 pathway while dampening external representations via the extra-hippocampal entorhinal inputs at longer timescales.

In conclusion, our findings suggest that mTORC1 activity in SOM-INs contributes to efficient learning and accurate hippocampal memory consolidation and is necessary for learning-induced persistent potentiation of excitatory inputs of CA1 SOM-INs. At the network level, SOM-IN mTORC1 mechanisms contribute to regulation of metaplasticity of the SC pathway in pyramidal cells. Thus, mTORC1-mediated plasticity in SOM-INs might open a permissive temporal window for distributed learning such as spatial learning, and memory retrieval.

## Materials and Methods

### Subjects

All animal procedures and experiments were performed in accordance with the Université de Montréal animal care committee regulations. Mice were group housed (2-4 per cage), maintained with food and water ad libitum and on a 12 hr light/dark cycle with all testing performed during the light phase. Knock-in mice with an internal ribosome entry site (IRES)-linked Cre recombinase gene downstream of the *Sst* locus (*Sst*^ires-Cre^ mice, Jackson Laboratory #013044, Bar Harbour, ME). *Sst*^ires-Cre^ mice were crossed with *Rosa26*^lsl-EYFP^ reporter mice (Ai3, Jackson Laboratory #007903) for Cre-dependent Enhanced Yellow Fluorescent Protein (EYFP) expression in SOM-INs. *Sst*^ires-Cre^;*Rosa26*^lsl-EYFP^ mice were crossed with floxed *Rptor* mice (Jackson Labs #013188) for cell-specific knock-out of *Rptor* in SOM cells. *Sst*^ires-Cre^;*Rosa26*^lsl-EYFP^;*Rptor*^WT/fl^ heterozygous offsprings were backcrossed together to obtain *Sst*^ires-Cre^;*Rosa26*^lsl-EYFP^;*Rptor*^fl/fl^ homozygous knock-out mice (termed Som-Raptor-KO mice), and compared to the *Rptor* wild-type mice *Sst*^ires-Cre^;*Rosa26*^lsl-EYFP^;*Rptor*^WT/WT^ (termed Som-Raptor-WT mice). All strains were maintained on a C57BL/6N background. *Sst*^ires-Cre^;*Rosa26*^lsl-EYFP^ mice were crossed with floxed *Tsc1* mice (Jackson Labs # 005680) for cell-specific knock-down of *Tsc1* in SOM cells. *Sst*^ires-Cre^;*Rosa26*^lsl-EYFP^;*Tsc1*^WT/fl^ heterozygous offsprings (termed Som-TSC1-KO mice) were compared with *Sst*^ires-Cre^;*Rosa26*^lsl-EYFP^;*Tsc1*^WT/WT^ control mice (termed Som-TSC1-WT mice).

Molecular biology and immunohistochemistry experiments were performed on mice of both sex, electrophysiology experiments in mature acute slices and behavioral tests were performed on male mice only.

### EYFP-labelled interneuron distribution

Distribution of EYFP-labelled interneurons was determined by fluorescence microscopy from hippocampal sections of Som-Raptor-WT and -KO mice (6-12 weeks-old). Animals were deeply anesthetized intra-peritoneally with sodium pentobarbital (MTC Pharmaceuticals, Cambridge, ON), perfused transcardially with ice-cold 0.1M phosphate buffer (PB) and 4% para-formaldehyde in 0.1M PB (PFA) and the brain isolated. Post-fixed brains were cryoprotected in 30% sucrose and coronal brain sections (50 μm thick) were obtained using freezing microtome (Leica SM200R; Leica, Wetzlar, Germany). Sections were mounted in ProLong^®^ Gold (Invitrogen, Carlsbad, CA) and examined using an epifluorescence microscope (Nikon Eclipse E600; Nikon, Tokyo, Japan). EYFP-positive interneuron density was assessed in the hippocampal CA1 oriens layer.

### Hippocampal slice culture

Organotypic hippocampal slice cultures were obtained from Som-Raptor-WT and -KO mice (4-5 day-old), as described previously (*8*). In brief, the brain was removed and dissected in HBSS (Invitrogen)-based medium. Cortico-hippocampal slices (400 μm thick) were then obtained using a McIlwain tissue chopper (The Cavey Laboratory Engineering Co. Ltd, Surrey, UK). After dissection, slices were placed on Millicell culture plate inserts (Millipore, Burlington, MA) in OptiMEM (Invitrogen) supplemented with Glutamax I and horse serum (Invitrogen) kept at 37°C in a humidified atmosphere (95% air, 5% CO2) for 3–7 d.

### Chemical induction protocol for late-LTP

Slices received three applications (10 min duration each at 30 min intervals) of the mGluR1/5 agonist (S)-3,5-dihydroxyphenylglycine (DHPG, 5 μM; Abcam, Cambridge, UK) in the presence of the mGluR5 antagonist 2-methyl-6 (phenylethynyl)-pyridine (MPEP, 25 μM; Tocris Bioscience, Bristol, UK). Inhibitors were applied concomitantly from 30 min before to 30 min after DHPG/MPEP treatment. LY367385, rapamycin and PP242 were purchased from Tocris, Millipore, and LC Laboratories (Woburn, MA), respectively. After treatments, cultured and acute slices were allowed to recover for 24h and 2h, respectively, before recordings. Experimenters were blind to all treatment groups and mice genotype.

### Immunofluorescence

Co-localization of Raptor immunofluorescence was determined in EYFP-positive interneurons. Coronal brain sections (50 μm thick) were permeabilized with 0.5% Triton X-100 in PBS (15 min) and unspecific binding was blocked with 10% normal goat serum in 0.1% Triton X-100/PBS (1h). Mouse monoclonal Raptor (1/500; Millipore) antibody was incubated 48 hours at 4°C. Sections were subsequently incubated at room temperature with Rhodamine-conjugated goat anti-mouse IgG1 (1/200; 90 min; Jackson Immunoresearch Laboratories, West Grove, PA). Sections were mounted in ProLong^®^ Gold and imaged as for EYFP-labelled interneuron distribution. The number of EYFP-positive interneurons in CA1 stratum oriens with co-localization of Raptor immunofluorescence were counted and expressed as percent of the total EYFP-positive cells.

Co-localization of somatostatin immunofluorescence in EYFP-labelled interneurons was performed on organotypic slices obtained as described before. Slices were fixed during 24h with 4% PFA, cryoprotected in 30% sucrose and re-sectioned (50 μm thick) using freezing microtome. Sections were permeabilized 30 min with 0.5% Triton X-100/PBS and unspecific binding blocked as above. Sections were incubated overnight at 4°C with rabbit polyclonal somatostatin 28 antibody (1/2000; Abcam) and subsequently at room temperature with Alexa 594-conjugated goat anti-rabbit IgGs (1/500, 90 min; Abcam). Sections were mounted, imaged and co-localization measured as described above for Raptor immunofluorescence.

### S6 immunophosphorylation assay

Mice (3-5 weeks-old) were deeply anesthetized with sodium pentobarbital and perfused transcardially with ice-cold ACSF containing (in mM): 110 choline-chloride, 2.5 KCl, 7 MgCl2, 26 NaHCO3, 7 dextrose, 1.3 ascorbic acid and 0.5 CaCl2, and saturated with 95% O2-5% CO2. Coronal hippocampal slices (300 μm) were obtained using a vibratome (Leica VT 1000S) and transferred at room temperature into normal oxygenated ACSF containing (in mM): 124 NaCl, 2.5 KCl, 1.25 NaH2PO4, 2 MgCl2, 2 CaCl2, 26 NaHCO3, 10 dextrose, 1.3 ascorbic acid. After 1 hour recovery period, slices received the late-LTP chemical induction protocol at 31-33°C as described above. Slices were fixed after treatment and cryoprotected in 30% sucrose 24 hours later. Slices were re-sectioned in 50 μm thick sections using a freezing microtome (Leica SM200R). Sections were permeabilized with 0.3% Triton X-100 in PBS (15 min) and unspecific binding was blocked with 10% normal goat serum in 0.1% Triton X-100/PBS (1h). Rabbit polyclonal phospho-S6 ribosomal protein (1/200; anti-phospho-S6S235/236; Cell Signaling, Danvers, MA) antibodies were incubated 48 hours at 4°C. Sections were subsequently incubated at room temperature with Alexa 594-conjugated goat anti-rabbit IgGs (1/500; 90 min; Jackson Immunoresearch). Images were acquired using confocal microscope (LSM510, Zeiss, Oberkochen, Germany) at excitation 488 and 543 nm. Images from different treatment/groups were acquired using the exact same parameters. Cell fluorescence was quantified using Fiji (ImageJ, NIH) by comparing integrated density in cells corrected for background.

### Western Blotting

Total hippocampus (10 weeks-old mice) and slice cultures (3 slices cortex-free pooled per sample) were collected and protein extracted using ice-cold radioimmunoprecipitation assay buffer containing: 50 mM Tris pH 7.4, 150 mM NaCl, 2 mM EDTA, 1% Triton X-100, 0.5% sodium desoxycholate, 0.1% sodium dodecyl sulfate (SDS), 200 μM NaF, 200 μM Na3VO4 and protease inhibitor (Cocktail inhibitor set I; Millipore) (20 min, 4°C). Lysates were centrifuged at 19 000 g (20 min, 4°C) and protein concentration from supernatant was determined according to bicinchoninic acid method using bovine serum albumin as standard. Fifteen to thirty micrograms of proteins (slice culture or total hippocampus extracts respectively) were separated by 7% (Raptor) or 12% (p-S6) SDS-polyacrylamide gel electrophoresis and transferred onto polyvinilidene fluoride membrane. The membranes were blocked with 5% non-fat skin milk dissolved in Tris-buffered saline-0.1% tween 20 pH 7.4 (1h30, room temperature) and incubated with rabbit polyclonal anti-phospho-S6S235/236 (1/1000; Cell Signaling) or rabbit monoclonal anti-Raptor (1/500; Cell Signaling) overnight at 4°C. Membranes were then incubated with horseradish peroxidase-conjugated anti-rabbit IgGs (1/20000; Jackson Immunoresearch) for 1h30 at room temperature. Immunoreactive bands were detected by enhanced chemiluminescence plus (Perkin Elmer, Waltham, MA). Membranes were next stripped with buffer containing 0.2 M glycine pH 2.2, 0.1% SDS and reprobed with antibodies detecting level of total S6 (1/2000; Cell Signaling) and/or tubulin (1/1000; Cell Signaling) overnight at 4°C. All immunoreactive bands were scanned with a desktop scanner and quantified using Quantity One software (BioRAD, Hercules, CA).

### Acute hippocampal slice preparation

Acute slices were prepared from 7- to 10-week old Som-Raptor-WT and -KO mice. Animals were anesthetized with isoflurane inhalation and the brain was rapidly removed and placed in ice-cold sucrose-based cutting solution containing the following (in mM): 75 sucrose, 87 NaCl, 2.5 KCl, 1.25 NaH2PO4, 7 MgSO4, 0.5 CaCl2, 25 NaHCO3, 25 glucose, 11.6 ascorbic acid and 3.1 pyruvic acid, pH 7.4, and 295 mOsmol/L. A block of tissue containing the hippocampus was prepared and 300 or 400 μm (for whole-cell and field recordings, respectively) transverse hippocampal slices were cut with a Leica VT1000S vibratome. Slices were transferred for recovery for 30 min to a holding chamber in artificial cerebral spinal fluid (ACSF) containing the following (in mM): 124 NaCl, 2.5 KCl, 1.25 NaH2PO4, 1.3 MgSO4, 2.5 CaCl2, 26 NaHCO3, and 10 glucose (pH 7.3–7.4, 295–305 mOsmol/L) at 30°C and subsequently maintained at room temperature (20–22°C) for at least 90 min until use. Both cutting solution and ACSF were saturated with 95% O2/5% CO2.

### Whole-cell recordings

For experiments in cultured slices, culture plate inserts were transferred to ACSF containing the following (in mM): 124 NaCl, 2.5 KCl, 1.25 NaH2PO4, 4 MgSO4, 4 CaCl2, 26 NaHCO3, and 10 glucose (pH 7.3–7.4, 295–305 mOsmol/L) maintained at room temperature for at least 30 min until use. Acute and cultured slices were transferred to a submersion chamber perfused (3–4 ml/min) with ACSF at 31 ± 0.5°C, CA1 and CA3 regions were disconnected by a surgical cut and slices kept for an additional 30 minutes submerged before recording. EYFP-expressing CA1 interneurons were identified using an upright microscope (Nikon Eclipse, E600FN), equipped with a water-immersion long-working distance objective (40x, Nomarski Optics), epifluorescence and an infrared video camera. Whole-cell voltage-clamp recordings were obtained using borosilicate glass pipettes (2-5 MΩ; WPI) filled with intracellular solution containing the following (in mM): 120 CsMeSO3, 5 CsCl, 2 MgCl2, 10 HEPES, 0.5 EGTA, 10 Na2-phosphocreatine, 2 ATP-Tris, 0.4 GTP-Tris, 0.1 spermine, 2 QX314, and 0.1% biocytin, pH 7.2–7.3, and 280 ± 5 mOsmol. For whole-cell current-clamp recordings, the intracellular solution contained the following (in mM): 120 KMeSO4, 10 KCl, 10 HEPES, 0.5 EGTA, 10 Na2-phosphocreatine, 2.5 MgATP, 0.3 NaGTP, and 0.1% biocytin (pH 7.4, 300 mOsmol/L). Data was acquired using a Multiclamp 700B amplifier (Molecular Devices, San Jose, CA), digitized at 20 kHz using Digidata 1440A and pClamp 10 (Molecular Devices). Recordings were low-pass filtered at 2 kHz. Access resistance was regularly monitored during experiments and data were included only if the holding current was stable and access resistance varied less than 20% of initial value. EPSCs were recorded in the presence of (2R)-amino-5-phosphonovaleric acid (AP5; 50 μM, Sigma, St-Louis, MO) and SR-95531 (gabazine; 5 μM, Sigma) to block NMDA and GABAA receptors, respectively. Data were acquired and analysed by an experimenter blinded for the genotype and the treatment of the animals.

Spontaneous EPSCs were recorded in voltage-clamp mode over a period of 1-5 min and 300 consecutive EPSCs analyzed for frequency and amplitude (MiniAnalysis, Synaptosoft, Fort Lee, NJ). Evoked EPSCs were elicited using constant current pulses (50 μs duration) via an ACSF-filled bipolar theta-glass electrode (Harvard Apparatus, Holliston, MA) positioned approximately 100 μm lateral to the recorded cell soma at the border between CA1 stratum oriens and the alveus. Putative single-fiber EPSCs were evoked at 0.5 Hz using minimal stimulation (success rate = 40-50%). EPSC potency (EPSC amplitude excluding failures) was averaged over a 10-15 min period (pClamp 10). Because the failure rate was an adjusted parameter, we used EPSC potency to characterize amplitude changes in evoked transmission. Input-output function was studied by delivering current pulses of incremental intensity (0-600 μA, 50 μA steps); 3-10 trials per pulse intensity were delivered and responses averaged (including failures) to determine EPSC amplitude (initial EPSC peak). Linear regressions were applied on individual averaged responses; the slope (synaptic gain) and x-intercept (minimal stimulation intensity) of the linear regression of the input-output relationship were measured.

Membrane and firing properties of EYFP-labeled SOM interneurons were measured in current-clamp recordings (*38*). Resting membrane potential was measured with the holding current I=0 pA immediately after break-in. Input resistance (Rm) was measured using a linear regression of voltage deflections (± 15 mV max) in response to current steps (500 ms, 5 pA increment, holding membrane potential −60 mV), excluding responses with voltage deflections with activation of voltage-dependent conductance (voltage sag or ramps). Membrane time constant was calculated from the mean responses to 20 successive hyperpolarizing current pulses (−5 pA; 500 ms), determined by fitting voltage responses with a single exponential function and used to calculate specific membrane capacitance. Action potential (AP) threshold was taken as the voltage at which the slope trajectory reached 10 mV/ms. AP amplitude was the difference in membrane potential between threshold and peak. AP half-width was calculated as AP duration at half-amplitude. Fast afterhyperpolarization (fAHP) amplitude was calculated as the difference between threshold and the negative peak after the AP. The time difference between the current pulse onset and the AP peak was defined as AP latency. The rheobase was measured as the minimal current amplitude necessary to evoke an action potential. AP properties were measured for the first AP elicited by a 500 ms depolarizing current pulse just sufficient to reach threshold. The sag index was determined from a series of negative current steps (500 ms duration, 10 pA steps). From the V–I plots, the peak negative voltage deflection (Vhyp) and the steady-state voltage deflection (Vsag, calculated for the last 50 ms of the current step) were used to calculate the index as the ratio (Vrest – Vsag) / (Vrest – Vhyp), for current injections corresponding to Vsag = −80 mV.

### Electrical stimulation protocol for late-LTP induction

Late-LTP was induced by electrical stimulation in acute slices using a concentric bipolar Pt/Ir electrode (FHC, Bowdoin, ME) positioned in the stratum oriens close to the alveus. Late-LTP induction protocol consisted of 4 trains of theta-burst stimulations (TBS) at 5 min intervals. Each TBS train consisted of 3 episodes (at 30 s intervals) of 5 bursts (at 250 ms inter-burst intervals) of 4 pulses at 100 Hz (*18, 19*). Whole cells recordings were obtained from EYFP-positive interneurons located approximately 100 μm lateral to the stimulating electrode at 2h after late-LTP induction or after a similar period in unstimulated slices. LTP was assessed by recording EPSCs evoked by minimal stimulation through an ACSF-filled bipolar theta-glass electrode positioned at approximately the same site as the stimulating electrode used for LTP induction.

### Field recordings

Field EPSPs (fEPSPs) were recorded as previously (*19*) in CA1 stratum radiatum with glass electrodes (1–3 MΩ; WPI, Saratosa, FL) filled with ASCF. Schaffer collaterals were stimulated (0.1 ms duration; 30s-1) using a concentric bipolar Pt/Ir stimulating electrode (FHC) placed in stratum radiatum proximal to the CA3 region. A second concentric bipolar Pt/Ir stimulating electrode was positioned in the oriens/alveus border proximal to the subiculum for theta-burst conditioning trains (as described above). Field potentials were recorded with a differential extracellular amplifier (Microelectrode AC Amplifier Model 1800, A-M Systems, Sequim, WA), filtered at 2 kHz, digitized at 10 kHz (Digidata 1440A), and analyzed with pClamp10 (Molecular Devices). Stimulus intensity was adjusted to elicit 35% of maximal fEPSP. Basal synaptic transmission was assessed by recording fEPSP in response to incremental stimulations (20 to 180 μA, 20 μA steps). Short term plasticity was assessed by fEPSP in response to paired-pulse stimulations with incremental delays (20 to 110 ms, 30 ms steps). LTP was induced at CA1 Schaffer collateral synapses through the stimulating electrode in the stratum radiatum by a weak high-frequency stimulation train (wHFS; 750 ms, 100 Hz). Two hours before LTP induction in the Schaffer collateral pathway, the conditioning TBS protocol to induce late-LTP in SOM interneurons (as described above) was applied at the oriens/alveus border. fEPSP slope was measured at 10–90% of fEPSP amplitude.

### Confocal imaging and cell reconstruction

Neurons were labeled with 0.1 % biocytin added to the whole cell patch pipette internal solution. At the end of experiments, hippocampal slices were postfixed overnight at room temperature in 4% paraformaldehyde in 0.1M phosphate buffer and 0.1 % Triton. The morphology of stained neurons was revealed using 1/1000 cy3-conjugated streptavidin (Rockland Immunochemical, Pottstown, PA) incubated overnight at room temperature. Cy3 was excited at 543 nm wavelength and images were captured using an LSM 880 confocal microscope (Zeiss). Stack images (1 μm steps) were acquired through a 40x water-immersion objective to perform three-dimension reconstructions and Sholl analysis of SOM interneurons using the Simple Neurite Tracer and Sholl Analysis Fiji plugins (ImageJ, NIH). SOM interneuron reconstructions revealed that in each group (Som-Raptor-WT and -KO mice), 90% displayed all features of OLM type of SOM interneurons, the remaining 10% showing dense axonal arborisation in strata oriens and radiatum, typical of bistratified cells or axonal collateral running out from the hippocampus typical of projection cells (*9*).

### Behavioral experiments

Before behavioral experiments, mice were gently handled daily for three days (~1 min per mouse) to habituate mice to the experimenter and reduce the stress related to the experimental handling. Mice were 8-13 weeks old. The experimenter was blind to the genotype of the mice. In all experiments, mice were first video-tracked at 25 frames per second and their movements subsequently analyzed using a position tracking system (Smart 3.0, PanLab, Barcelona, Spain). Mice performed the open-field test first; the Barnes maze test started the next day and the fear conditioning occurred 5 days after the end of the Barnes maze session. Half of the mice were subjected to fear conditioning only.

#### Open-field test

Mice were allowed to freely explore a home-made circular (Som-Raptor-WT and -KO mice, 60 cm diameter, 25 cm height) or square (Som-TSC1-WT and -KO mice, PanLab) open-field for 5 min. Locomotion was evaluated by measuring the number of zone transitions using a pattern divided into 25 (circular OF) or 21 (squared OF) zones of equal size. Anxiety was assessed by measuring the time spent in a 20 cm radius/side central zone versus in a 20 cm broad peripheral annulus.

#### Fear conditioning

Mice were trained in conditioning chambers that were housed in sound- and light-isolated cubicles (Coulbourn Instruments, Holliston, MA). The chambers contained a stainless steel grid floor, overhead LED lighting, camera and were supplied with background noise (60 dB) by an air extractor fan. The experimental protocol was based on Ruediger and coworkers (*10*) with slight modifications. The training context was rectangular with transparent walls and was cleaned with 1% acetic acid before and after each trial. Two neutral contexts were designed to assess contextual generalization and discrimination. One had a triangular shape, transparent walls and was cleaned with 70% ethanol before and after each trial. This context was considered novel but similar to the training context because they shared some features. Another neutral context had a circular shape, opaque black and reflective walls, Plexiglas floor and was cleaned with 70% ethanol before and after each trial. This context was considered novel and distinct to the training context. Freezing was assessed using FreezeFrame (Coulbourn Instruments). Once placed in the conditioning chamber, mice were allowed to freely explore for 2.5 min, and then received 5 presentations of unconditioned and conditioned stimuli (1 s foot shock, 0.8mA; where indicated, 10 kHz tone for 10 s, 70 dB sound pressure level, 30 s interleaved). The last 1 s of each tone was paired with the unconditioned stimulus. To test for cued fear conditioning, mice explored the novel distinct context for 2.5 min, followed by 5 tone presentations (tone-dependent freezing). Contextual fear conditioning involved the same protocol, but without the tone component. To test for contextual fear memory, mice were returned to the training context during a test period of 2.5 min, at 1 or 24 h after conditioning, to assess short- or long-term memory, respectively. To test for contextual discrimination after fear conditioning, a within-subjects design was used. On the test day, 5 h after the test in the training context, mice were pseudo-randomly distributed across the two novel neutral contexts and freezing was assessed during 2.5 min. Discrimination ratio was calculated as the amount of freezing in (training context) / (training context + novel context) (*39*). A ratio of 1 indicates that mice were able to discriminate the contexts perfectly, and a ratio of 0.5 means that they were unable to discriminate. For a subset of training-induced persistent plasticity experiments, mice were subjected to contextual fear conditioning 30 min after IP injection of 1 mg/kg JNJ16259685 (Tocris) or vehicle.

#### Barnes maze

This test was used to assess hippocampus-dependent spatial learning (*24*) using an elevated platform (PanLab). The experimental protocol was based on Sunyer and coworkers (*40*) with slight modifications. Mice were trained to use spatial cues to find a small dark escape chamber under the platform termed “target”. The target is retained at the same position relative to the room, while the platform is rotated with each trial to discourage use of the intra-maze odor cues. In addition, the platform, the starting chamber and the escape box were thoroughly cleaned (Versaclean 10 %) between every single trial to reduce any possible scent trails. Assignment of target location was balanced among experimental groups. An aversive pulsed noise (85 dB) went on during each trial and switched off when the mouse entered in the target. One day after a familiarization period (day 1), mice were trained in 4 daily sessions with an inter-trial interval of 15 min for 4 days. During the training, mice were left 10 s in a dark cylinder (starting chamber) in the center of the maze, allowed to freely explore the maze for a maximum duration of 3 min and left 1 min in the escape box after each trial. If a mouse failed to find the t by the end of the 3 min period, it was gently guided to it. Video tracking and data analysis were performed using Smart 3.0 (PanLab). On day 6, the mice were exposed to a probe trial in which the escape box was closed. Mice were allowed to explore the maze for 3 min. The number of errors, the latency and the distance travelled before the first reaching of the target (primary search) were collected. During the total search (90 s), the time spent in the quadrants (target, left, right and opposite, excluding a 15 cm diameter central zone) and the number of visits for each hole were collected. Non-target data was defined as the average number of visits in the 19 non-target holes. The selective search ratio was defined as the number of visits in (target) / (target + non-target). A ratio of 1 indicates that a mouse visited only the target hole, and a ratio of 0.5 means that it visited target and non-target holes equally. Search strategies were also collected and defined into three categories: 1) direct (spatial) – moving directly to the target hole or to an adjacent hole before visiting the target; 2) serial (thygmotactism) – the first visit to the target hole preceded by visiting at least two adjacent holes in serial manner, in clockwise or counter-clockwise direction; and 3) random (or mixed) – hole searches separated by crossing through the centre of the maze or unorganized search (*24, 25*). One search strategy was attributed for every single trial of the training and for the primary search of the probe test.

### Statistical analysis

No statistical methods were used to predetermine the sample size, but our sample sizes are similar to (or larger than) those generally employed in the field. Statistical analysis was performed using OriginPro 2016 (OriginLab, Northampton, MA). Data were tested for normality and homoscedasticity using the Shapiro-Wilk and the Brown–Forsythe tests, respectively. We used Student t tests (with Welch corrections for heteroscedasticity), two-way ANOVA with Tukey’s pairwise comparison tests (with Bonferroni adjustments for multiple comparisons) and two-way repeated-measures ANOVA when data passed normality and homoscedasticity assumptions. Mann-Whitney tests and Friedman ANOVA were used when it was not the case. All the tests were two-sided. In the figures, data are expressed as arithmetic mean and standard error of the mean (mean ± SEM). Asterisks denote statistical significance as calculated by the specified statistical tests (*, P < 0.05; **, P < 0.01; ***, P < 0.001, ns, not significant).

## Acknowledgments

**General**: We thank J. Pepin for help with slice cultures and chemical late-LTP experiments, and members of the Lacaille laboratory for helpful discussions and comments on the manuscript.

## Author contributions

J.A., I.L. and J.C.L. designed and directed experiments; J.A., A.J., A.K., E.H., A.L.F., A.S.R. and I.L. performed the experiments and analysed data; I.L. generated and validated transgenic mice lines; J.A. and J.C.L. conceived the project and wrote the paper with contributions from I.L.

## Data and materials availability

Requests for materials should be addressed to J.C.L.

## Supplementary Materials

Fig. S1. Chemically-induced persistent LTP at excitatory synapses onto CA1 SOM-INs depends on mGluR1a and mTOR.

Fig. S2. Intact SOM interneuron numbers and morphology in Som-Raptor-KO mice.

Fig. S3. Membrane and firing properties of SOM-INs from Som-Raptor-KO mice.

Fig. S4. Anxiety and locomotor activity of Som-Raptor-KO mice in the open field test.

Fig. S5. Spatial memory deficits in Som-Raptor-KO mice.

Fig. S6. Membrane and firing properties of SOM-INs from Som-TSC1-KO mice.

Fig. S7. Anxiety and locomotor activity of Som-TSC1-KO mice in the open-field test.

Fig. S8. Spatial memory strengthening in Som-TSC1-KO mice.

Fig. S9. A model for regulation of hippocampal memory by mTORC1 in CA1 SOM interneurons.

