## Supplementary material for "Regulation of hippocampal memory by mTORC1 in somatostatin interneuron"

This file includes:

Fig. S1. Chemically-induced persistent LTP at excitatory synapses onto CA1 SOM-INs depends on mGluR1a and mTOR.

Fig. S2. Intact SOM interneuron numbers and morphology in Som-Raptor-KO mice.

Fig. S3. Membrane and firing properties of SOM-INs from Som-Raptor-KO mice.

Fig. S4. Anxiety and locomotor activity of Som-Raptor-KO mice in the open field test.

Fig. S5. Spatial memory deficits in Som-Raptor-KO mice.

Fig. S6. Membrane and firing properties of SOM-INs from Som-TSC1-KO mice.

Fig. S7. Anxiety and locomotor activity of Som-TSC1-KO mice in the open-field test.

Fig. S8. Spatial memory strengthening in Som-TSC1-KO mice.

Fig. S9. A model for regulation of hippocampal memory by mTORC1 in CA1 SOM interneurons.

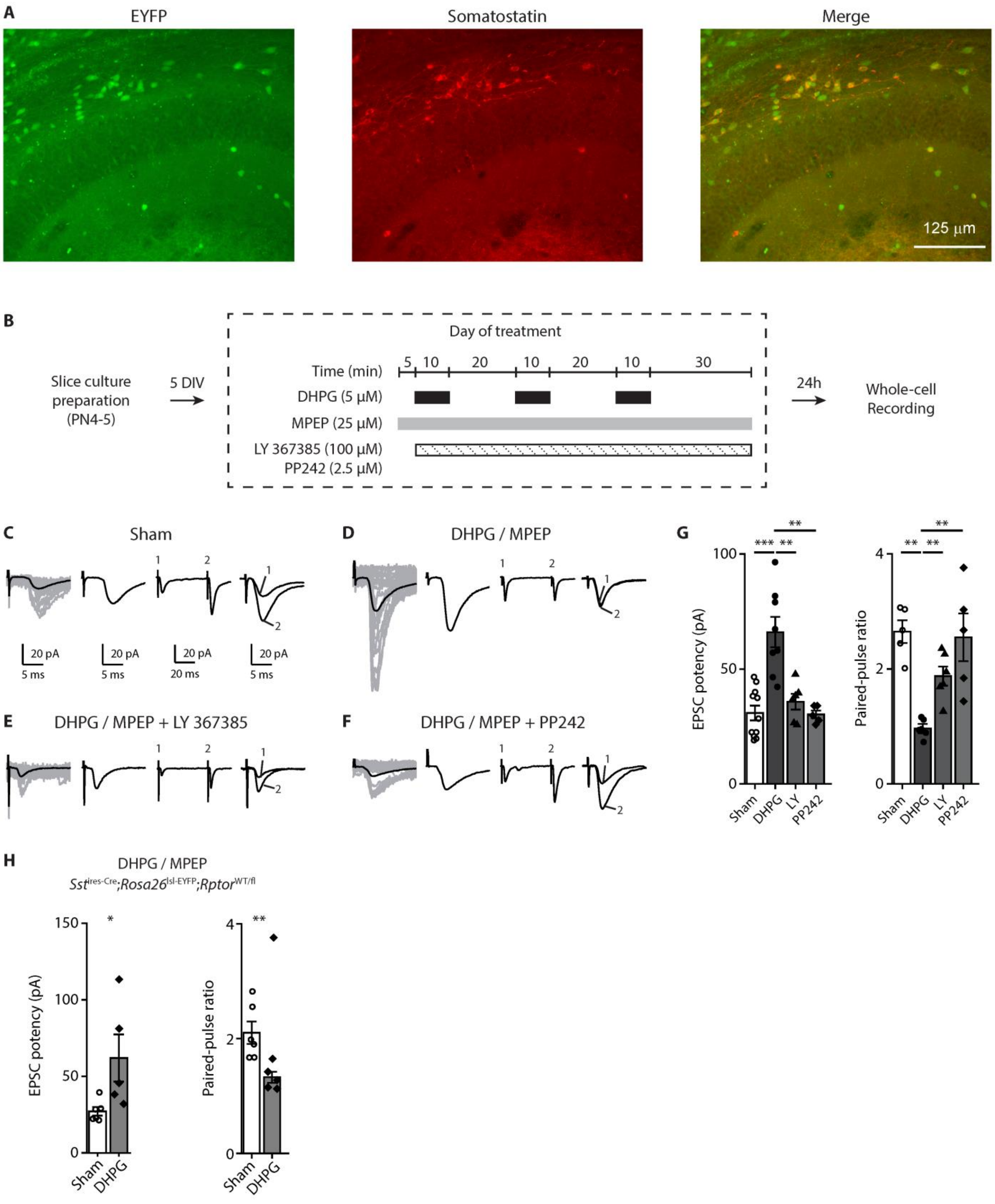
A
EYFP
Somatostatin
Merge
125 μm
B
Slice culture preparation (PN4-5)
5 DIV
Day of treatment
Time (min)
5 10 20 10 20 10 30
DHPG (5 μM)
MPEP (25 μM)
LY 367385 (100 μM)
PP242 (2.5 μM)
24h
Whole-cell Recording
C
Sham
20 pA
5 ms
20 pA
5 ms
20 pA
20 ms
20 pA
5 ms
D
DHPG / MPEP
E
DHPG / MPEP + LY 367385
F
DHPG / MPEP + PP242
G
EPSC potency (pA)
Sham
DHPG
LY
PP242
Paired-pulse ratio
H
DHPG / MPEP
Sst^ires-Cre;Rosa26^lsl-EYFP;Rptor^WT/fl
EPSC potency (pA)
Paired-pulse ratio
Sham
DHPG

**Fig. S1. Chemically-induced persistent LTP at excitatory synapses onto CA1 SOM-INs depends on mGluR1a and mTOR.** (**A**) Representative images of EYFP expression (left) and SOM immunofluorescence (middle) in hippocampal CA1 area from Sstires-Cre; Rosa26$^{lsl\text{-}EYFP}$ mice, showing 84.4 ± 2.6% of EYFP+ cells co-labelled for somatostatin (right) in stratum oriens. (**B**) Diagram of chemical late LTP induction and recording protocol in organotypic slice cultures. (**C-F**), Representative EPSCs evoked by minimal stimulation in EYFP expressing SOM-INs at 24h after induction in different conditions: sham treatment (**C**), and repeated mGluR1 stimulation alone (**D**), in the presence of the mGluR1a antagonist LY36775 (**E**) or mTOR inhibitor PP242 (**F**). From left to right: EPSCs including failures (individual traces in grey, average trace in black), averaged EPSC potency (excluding failures), average of EPSCs evoked by paired-pulse stimulation, and superimposed first and second EPSCs of average pair. g, Quantification of EPSCs after sham (n=11) and MPEP/DHPG alone (n=8), in the presence of LY36775 (n=6) or PP242 (n=5). EPSC potency (Kruskal-Wallis ANOVA, *P*=0.0004, Mann-Whitney tests, sham vs DHPG: *P*=0.0008, DHPG vs LY: *P*=0.006, DHPG vs PP242: P=0.004) and paired-pulse ratio (Kruskal-Wallis ANOVA, *P*=0.006, Mann-Whitney tests, sham vs DHPG: *P*=0.008, DHPG vs LY: *P*=0.005, DHPG vs PP242: *P*=0.008). (H) Quantification of EPSCs in SOM-INs of slice cultures of heterozygous Som-Raptor-KO mice (Sst$^{ires\text{-}Cre}$;Rosa26$^{lsl\text{-}EYFP}$;Rptor$^{WT/fl}$) after sham (n=6) and MPEP/DHPG (n=5) treatments. EPSC potency (Mann-Whitney test, *P*=0.022) and paired-pulse ratio ($t_9$=3.3, *P*=0.009).

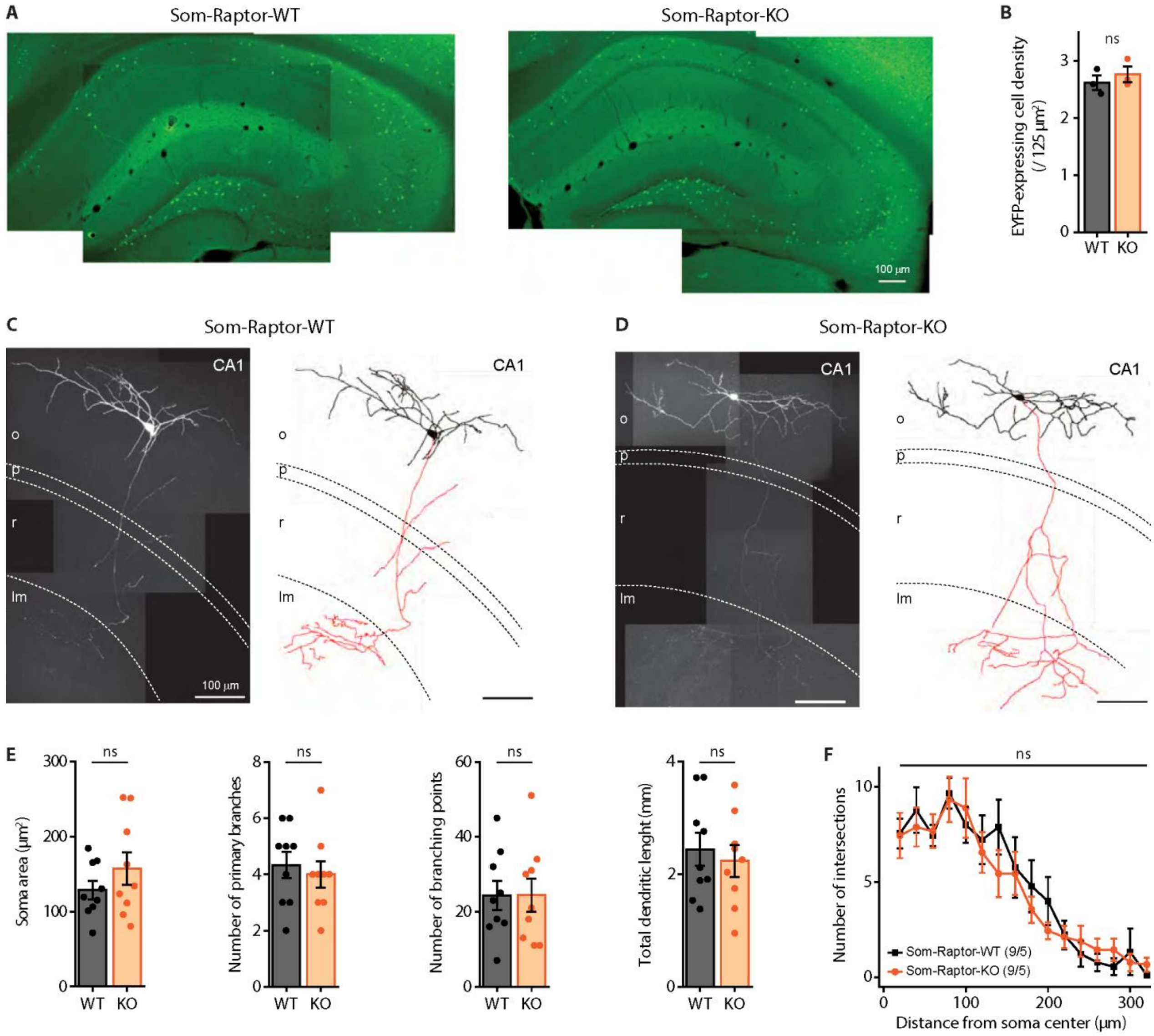
A
Som-Raptor-WT
Som-Raptor-KO
100 µm
B
ns
EYFP-expressing cell density (/ 125 µm²)
WT
KO
C
Som-Raptor-WT
CA1
o
p
r
lm
100 µm
D
Som-Raptor-KO
E
ns
Soma area (µm²)
Number of primary branches
Number of branching points
Total dendritic lenght (mm)
F
Number of intersections
Som-Raptor-WT (9/5)
Som-Raptor-KO (9/5)
Distance from soma center (µm)

50

51

**Fig. S2. Intact SOM interneuron numbers and morphology in Som-Raptor-KO mice.** (**A**) Montage of representative fluorescence images of EYFP expression in Som-Raptor-WT (left) and -KO (right) mice. (**B**) EYFP-expressing cell density in CA1 of Som-Raptor-WT (n=3 mice, 3 replicates each) and -KO (n=3 mice, 3 replicates each) mice ($t_4$=-0.8, $P$=0.49). (**C, D**) Montage of representative confocal images, maximum intensity z-projection (100 stacked images, 1 µm steps) of hippocampal CA1 SOM-IN filled with biocytin (left) and reconstructions (soma and dendrite in black, axon in red; right) from Som-Raptor-WT (**C**) and -KO (**D**) mice. As illustrated, the vast majority (90%) of EYFP-expressing SOM-INs recorded in whole-cell and filled with biocytin in the present study corresponded to the O-LM type of SOM-INs (the remaining 10% corresponding to bistratified cells and projection cells). The dashed lines indicate the approximate boundaries of strata oriens (o), pyramidale (p), radiatum (r) and lacunosum/moleculare (lm). (**E**) Somatic and dendritic morphometric parameters of biocytin-filled SOM-INs in Som-Raptor-WT (n=9 cells from 5 mice) and -KO (n=9 cells from 5 mice) mice. Soma area ($t_{16}$=1.1, $P$=0.27), number of primary branches ($t_{16}$=0.3, $P$=0.74), number of branching points ($t_{16}$=-0.3, $P$=0.75) and total dendritic length ($t_{16}$=-0.5, $P$=0.60). (**F**) Sholl analysis of dendritic arborisation of reconstructed SOM-INs (20 µm bins) in Som-Raptor-WT and -KO mice (two-way RM ANOVA, $F_{9,144,\text{interaction}}$=0.6, $P$=0.82).

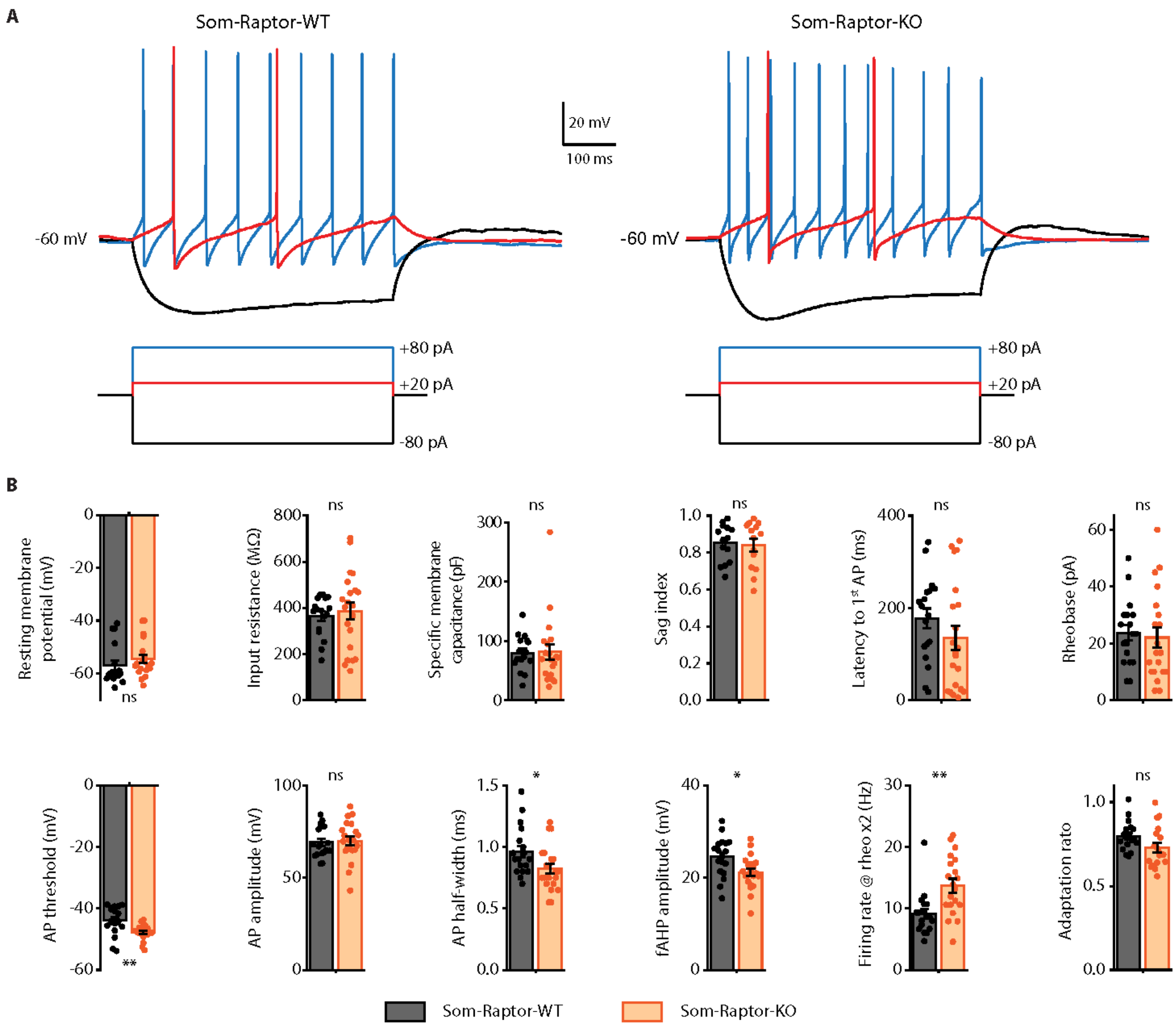
A
Som-Raptor-WT
Som-Raptor-KO
20 mV
100 ms
-60 mV
-60 mV
+80 pA
+20 pA
-80 pA
+80 pA
+20 pA
-80 pA
B
Resting membrane potential (mV)
Input resistance (MΩ)
Specific membrane capacitance (pF)
Sag index
Latency to 1st AP (ms)
Rheobase (pA)
AP threshold (mV)
AP amplitude (mV)
AP half-width (ms)
fAHP amplitude (mV)
Firing rate @ rheo x2 (Hz)
Adaptation rato
ns
*
**
Som-Raptor-WT
Som-Raptor-KO

72

73

**Fig. S3. Membrane and firing properties of SOM-INs from Som-Raptor-KO mice.** (**A**) Representative electrophysiological responses (top) of SOM-INs to current pulse injections (bottom) from Som-Raptor-WT (left) and -KO (right) mice. Depolarizing current pulses correspond to near threshold (red) and 4 × threshold (blue) stimulation. (**B**) Membrane and firing properties of SOM-INs from Som-TSC1-WT (n=14/18 cells from 9 mice) and -KO (n=14/20 cells from 10 mice) mice. Resting membrane potential (Mann-Whitney test, $P$=0.06), input resistance (Mann-Whitney, $P$=0.51), specific membrane capacitance (Mann-Whitney test, $P$=0.49), sag index ($t_{26}$=0.3, $P$=0.8), latency to first AP (Mann-Whitney test, $P$=0.14), rheobase (Mann-Whitney test, P=0.61), AP threshold (t-test with Welch correction, $t_{24.7}$=3.1, $P$=0.005), AP amplitude ($t_{37}$=-0.2, $P$=0.81), AP half-width ($t_{37}$=2.2, $P$=0.031), fAHP amplitude ($t_{37}$=-2.7, $P$=0.011), firing rate at rheobase x2 (Mann-Whitney test, $P$=0.0015) and adaptation ratio at rheobase x4 ($t_{35}$=2, $P$=0.055).

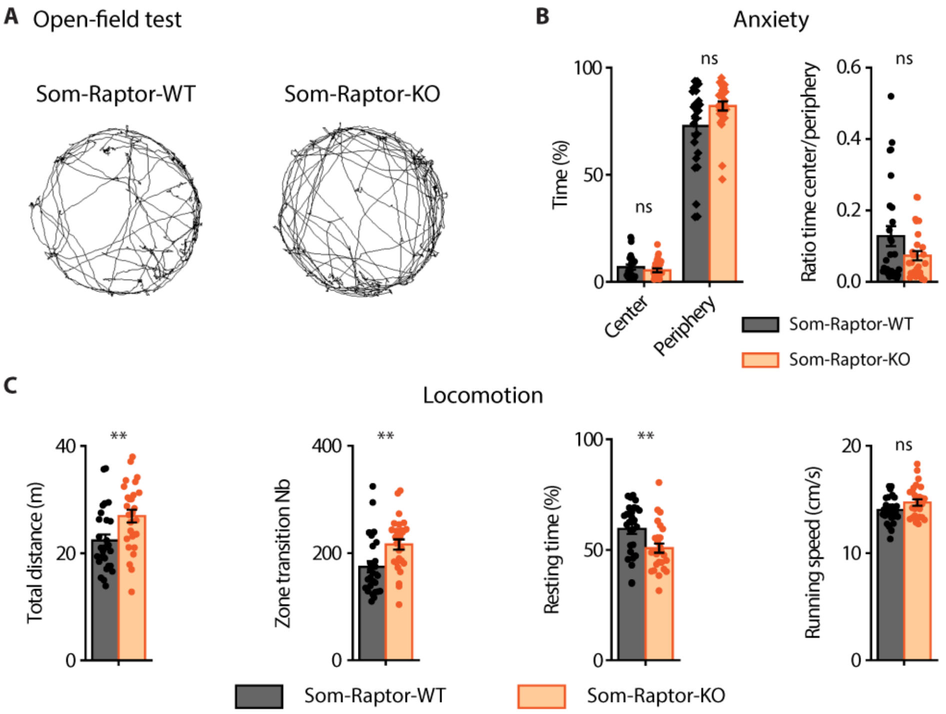


**Fig. 4. Anxiety and locomotor activity of Som-Raptor-KO mice in the open field test.** (**A**) Representative paths travelled by Som-Raptor-WT (left) and -KO (right) mice during 5 min free exploration in a circular open field. (**B**) Anxiety properties in the open-field test (n=27 mice in each group). Percentage of time spent in the center and in the periphery (Left, Mann-Whitney tests, Center: $P$=0.43 and Periphery: $P$=0.1) and center/periphery ratio (Right, Mann-Whitney test, $P$=0.27). (**C**) Locomotion properties in the open-field test. Total distance travelled ($t_{52}$=-2.8, $P$=0.008), zone transition number (Mann-Whitney test, $P$=0.002), percentage of resting time ($t_{52}$=2.8, $P$=0.007) and running speed ($t_{52}$=-1.9, $P$=0.057).

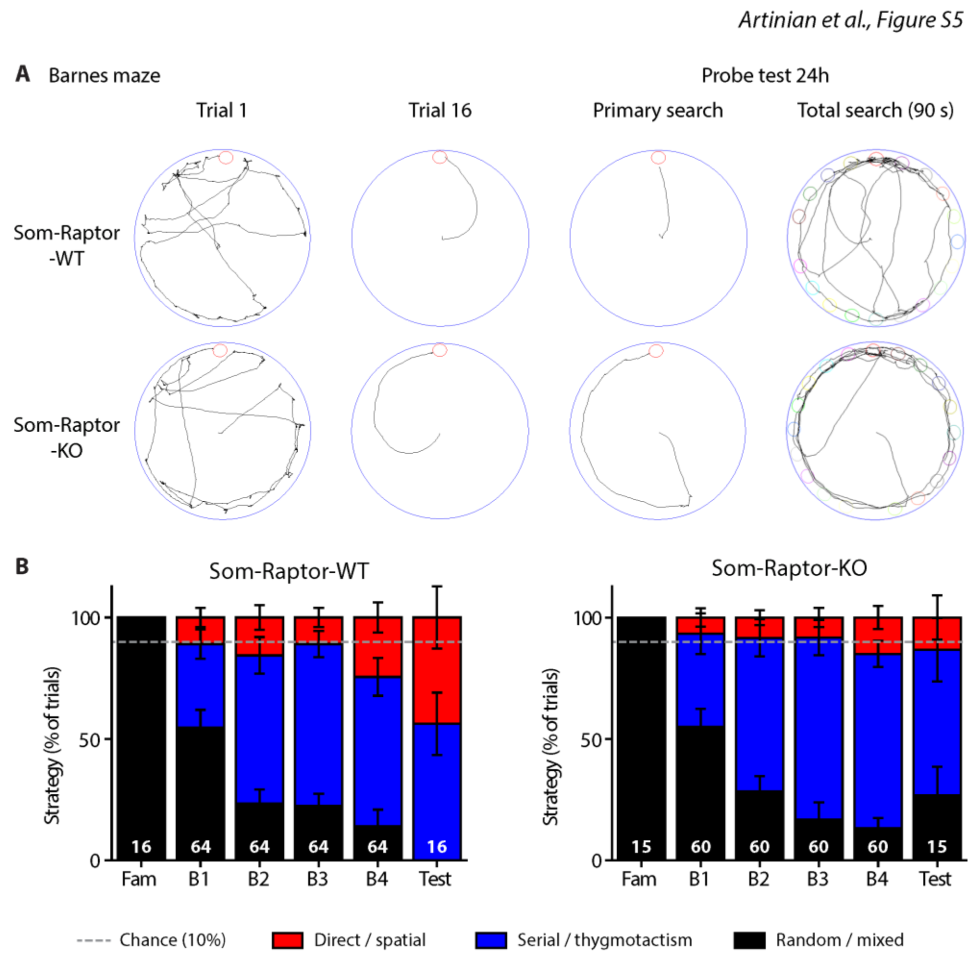


**Fig. S5. Spatial memory deficits in Som-Raptor-KO mice.** (**A**) From left to right: representative paths travelled by Som-Raptor-WT (top) and -KO (bottom) mice during the first and last ($16^{th}$) acquisition trials, as well as during the primary search and total search of the probe test. (**B**) Percentage of total trials performed using one of three types of strategy (spatial, thygmotactism and random/mixed) to solve the maze by Som-Raptor-WT (left, n=16) and -KO (right, n=15) mice. The dashed line represents chance level (10%) to find the target with a direct path. Numbers at the bottom of bars represent the total number of trials.

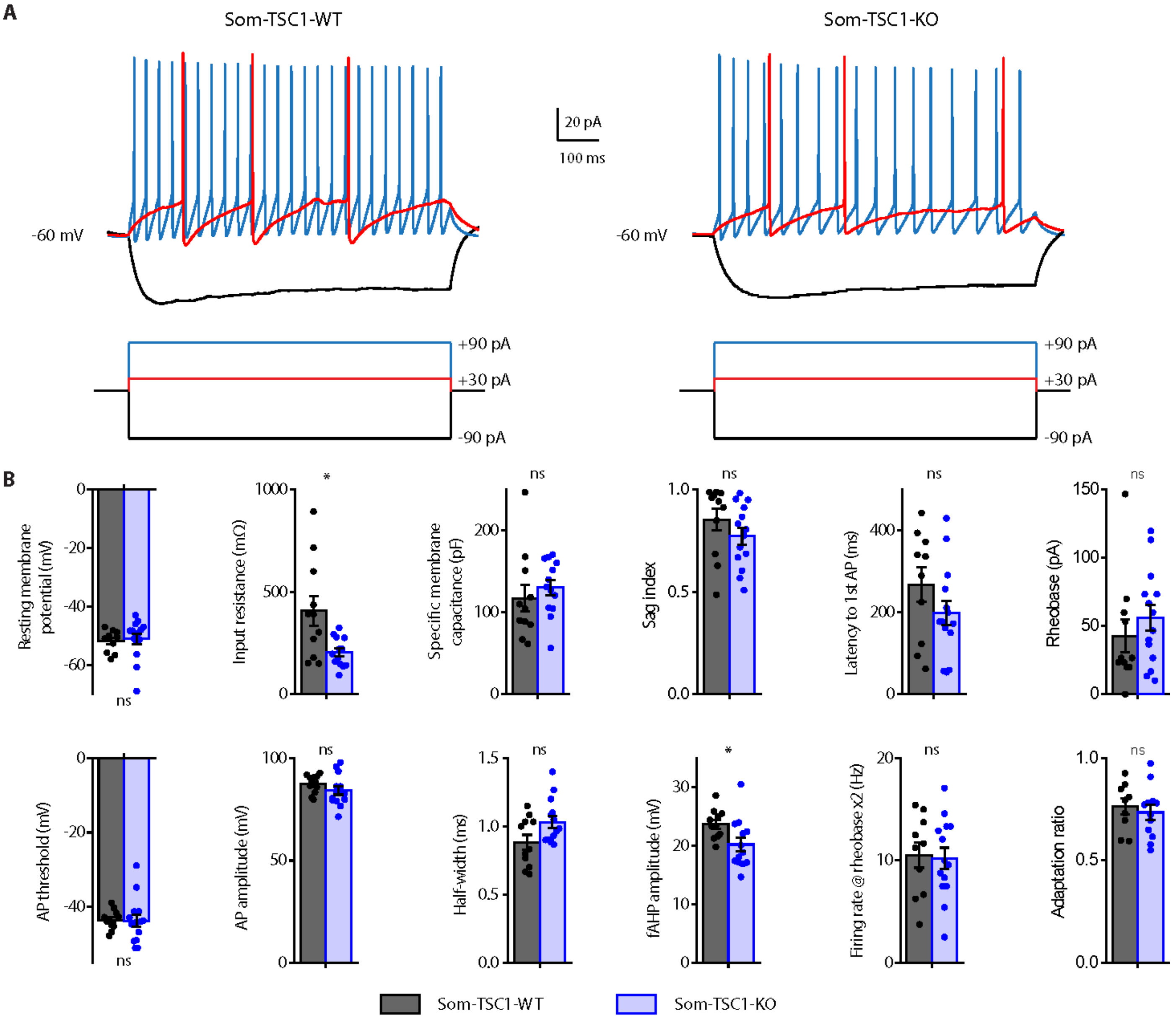
A
Som-TSC1-WT
Som-TSC1-KO
-60 mV
-60 mV
20 pA
100 ms
+90 pA
+30 pA
-90 pA
+90 pA
+30 pA
-90 pA
B
Resting membrane potential (mV)
ns
Input resistance (mΩ)
*
Specific membrane capacitance (pF)
ns
Sag index
ns
Latency to 1st AP (ms)
ns
Rheobase (pA)
ns
AP threshold (mV)
ns
AP amplitude (mV)
ns
Half-width (ms)
ns
fAHP amplitude (mV)
*
Firing rate @ rheobase x2 (Hz)
ns
Adaptation ratio
ns
Som-TSC1-WT
Som-TSC1-KO

**Fig. S6. Membrane and firing properties of SOM-INs from Som-TSC1-KO mice.** (**A**) Representative electrophysiological responses (top) of SOM-INs to current pulse injections (bottom) from Som-TSC1-WT (left) and -KO (right) mice. Depolarizing current pulses correspond to near threshold (red) and 3× threshold (blue) stimulation. (**B**) Membrane and firing properties of SOM-INs from Som-TSC1-WT (n=10/11 cells from 2 mice) and -KO (n=13/14 cells from 4 mice) mice. Resting membrane potential (Mann-Whitney, $P$=0.35), input resistance (t-test with Welch correction, $t_{11.5}$=2.7, $P$=0.02), specific membrane capacitance ($t_{22}$=-0.73, $P$=0.47), sag index (Mann-Whitney, $P$=0.12), latency to first AP ($t_{22}$=1.4, $P$=0.18), rheobase (Mann-Whitney, $P$=0.27), AP threshold (t-test with Welch correction, $t_{18.6}$=0.1, $P$=0.91), AP amplitude ($t_{23}$=1.2, $P$=0.23), AP half-width (Mann-Whitney, $P$=0.08), fAHP amplitude ($t_{22}$=1.2, $P$=0.022), firing rate at rheobase x2 ($t_{22}$=0.2, $P$=0.84) and adaptation ratio at rheobase x3 ($t_{19}$=0.5, $P$=0.61).

*Artinian et al., Figure S7*

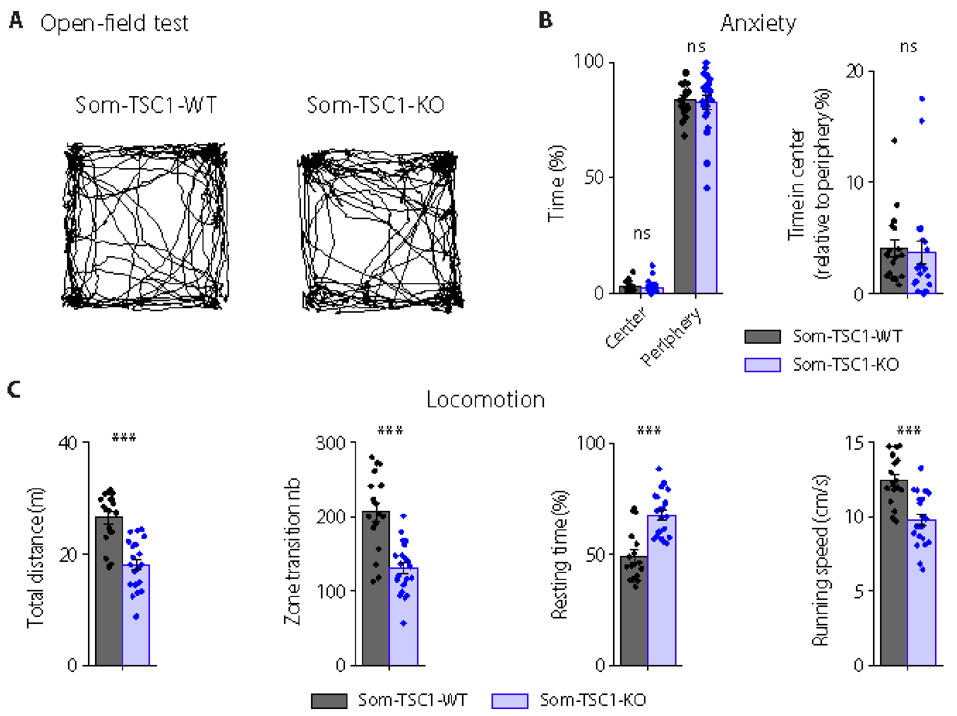


**Fig. S7. Anxiety and locomotor activity of Som-TSC1-KO mice in the open-field test.** (**A**) Representative paths travelled by Som-TSC1-WT (left) and -KO (right) mice during 5 min free exploration. (**B**) Anxiety properties of Som-TSC1-WT (n=17) and -KO (n=21) mice. Percentage of time freezing in the center and in the periphery (left, Mann-Whitney tests, Center: $P$=0.16; Periphery: $P$=0.84) and center/periphery ratio (right, Mann-Whitney tests, $P$=0.22). (**C**) Locomotion properties of Som-TSC1-KO mice. Total distance travelled (Mann-Whitney test, $P$=3.5$_E^{-5}$), number of zone transitions ($t_{36}$=5.4, $P$=5.1$_E^{-6}$), resting time (Mann-Whitney test, $P$=6.5$_E^{-5}$) and running speed ($t_{36}$=4.8, $P$=3.1$_E^{-5}$).

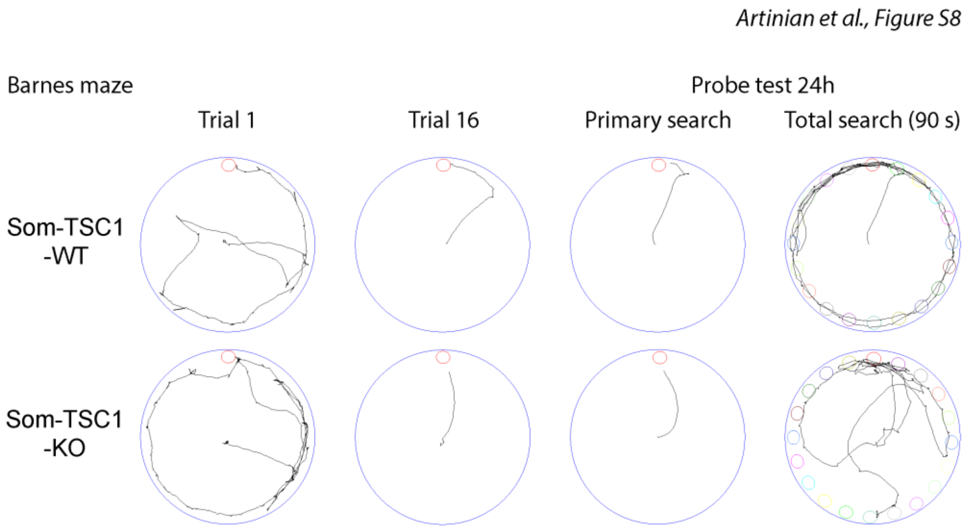


**Fig. S8. Spatial memory deficits in Som-TSC1-KO mice.** From left to right: representative paths travelled by Som-TSC1-WT (top) and -KO (bottom) mice during the first and last (16$^{th}$) acquisition trials, as well as during the primary search and total search of the probe test.

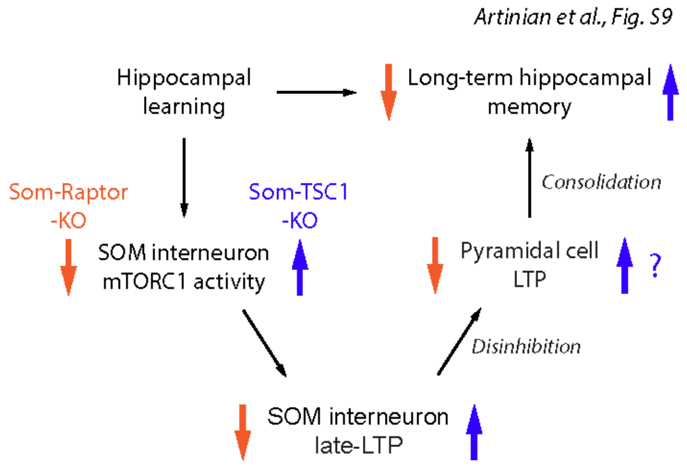


**Fig. S9: A model for regulation of hippocampal memory by mTORC1-mediated late-LTP in CA1 SOM interneurons.** Hippocampal learning engages CA1 pyramidal cell firing that increases mGluR1-mediated mTORC1 activity in SOM-INs and induces Hebbian late-LTP at SOM-INs excitatory input synapses. Late-LTP in SOM-INs is then converted into increased output firing that upregulates LTP at SC synapses onto local pyramidal cells by disinhibition. Facilitated LTP in the CA1 principal pathway supports memory consolidation and promotes long-term hippocampal memory for future recall.
